# Genomically hardwired regulation of gene activity orchestrates cellular iron homeostasis in Arabidopsis

**DOI:** 10.1101/2021.09.01.458651

**Authors:** En-Jung Hsieh, Wen-Dar Lin, Wolfgang Schmidt

## Abstract

Iron (Fe) is an essential micronutrient that plays pivotal roles as electron donor and catalyst across organisms. In plants, variable, often insufficient Fe supply necessitates mechanisms that constantly attune Fe uptake rates and recalibrate cellular Fe homeostasis. Here, we show that short-term (0.5, 6, and 12 h) exposure of *Arabidopsis thaliana* plants to Fe deficiency triggered massive changes in gene activity governed by transcription and alternative splicing (AS), regulatory layers that were to a large extent mutually exclusive. Such preclusion was not observed for genes that are directly involved in the acquisition of Fe, which appears to be concordantly regulated by both expression and AS. Generally, genes with lower splice site strengths and higher intron numbers were more likely to be regulated by AS, no dependence was on gene architecture was observed for transcriptionally controlled genes. Conspicuously, specific processes were associated with particular genomic features and biased towards either regulatory mode, suggesting that genomic hardwiring is functionally biased. Early changes in splicing patterns were, in many cases, congruent with later changes in transcript or protein abundance, thus contributing to the pronounced transcriptome-proteome discordance observed in plants.

## Introduction

Iron (Fe) is mineral nutrient with a plethora of vital functions across all kingdoms of life. In plants, Fe is a critical component of electron chains in photosynthesis and required for the biosynthesis of chlorophyll. Iron is abundant in the Earth’s crust, but its availability is often limited by interaction with other soil constituents, in particular at high redox and pH values. In most aerated soils, Fe is present in the form of sparingly soluble Fe(III) oxides which cannot readily be taken up by plants, causing leaf chlorosis, reduced yield, and decreased quality of edible plant parts. Low plant Fe content can jeopardise human health by causing Fe-deficiency anaemia, in particular in populations with a predominantly plant-based diet. To counteract limited Fe availability, plants have evolved a suite of mostly transcriptionally regulated responses that mediate the acquisition of Fe from recalcitrant pools in soils^1, 2^. In contrast to graminaceous species, which take up Fe^3+^ after secretion of Fe-avid phytosiderophores (strategy II)^3^, dicotyledonous plants acquire Fe^2+^ by the concerted action of processes that mobilize sparingly soluble Fe^3+^ by protonation, chelation, and reduction in response to imbalances caused by inadequate Fe supply (strategy I)^4–8^. The responses to Fe starvation are controlled by a complex network of homeostatic mechanisms that orchestrate the acquisition, transport, and cellular homeostasis of Fe^9–12^.

Due to the central role of ferrous Fe as an electron donor and catalyst, its absence or insufficiency compromises energy production and causes severe metabolic perturbations. Reduced respiration efficiency and impaired activity of Fe-containing key enzymes of the tricarboxylic acid (TCA) cycle such as aconitase or succinate dehydrogenase constitute major constraints in Fe-undernourished cells, necessitating extensive rerouting of carbon flow during periods of low Fe availability. Increased glycolysis rates upon Fe starvation, for instance, have been described for diverse eukaryote systems such as human cell lines^13^, macrophages^14^, and yeast^15^, suggesting that enhanced glycolytic flux is a conserved mechanism to compensate for decreased respiration. In Fe-deficient plants, pronounced metabolic changes have been observed across species^16–23^, which, with some notable exceptions, are not mirrored in transcriptomic profiles. The mechanisms that govern such metabolic rerouting have remained largely elusive.

Alternative splicing (AS) of pre-mRNAs, i.e., the process of selecting different combinations of splice sites for intron removal and exon ligation, contributes to the diversity of transcripts and proteins by producing multiple mRNA and protein isoforms, and may, at least partly, be causative for the lack of transcriptional footprints of Fe deficiency-induced alterations in central carbon metabolism. In contrast to animals, where exon skipping (ES) is dominating over other forms of AS, in plants, intron retention (IR) and alternative donor or acceptor splice sites (DA) are the prevalent forms of AS^24–26^. While ES is likely to produce functional proteins, the latter forms of AS often interrupt the open reading frame and lead to the introduction of premature stop codons, causing the formation of aberrant proteins or mRNA that is targeted to the nonsense-mediated decay pathway^27^. Between 60-80% of the multi-exonic plant genes are alternatively spliced, a percentage that can vary in response to environmental stimuli or stress^28–30^. Thus, in plants, AS constitutes a huge and mostly unexplored source of gene regulation of undisclosed significance, possibly fine-tuning the transcriptome to the prevailing environmental conditions.

How environmental information is communicated into the nucleus to alter pre-mRNA splicing patterns remains largely enigmatic. AS is driven by a large suite of mRNA-binding, spliceosome-associated proteins such as serine/arginine-rich (SR) proteins and heterogeneous nuclear ribonucleoproteins (hnRNPs) that binds to *cis*-regulatory sequences on the pre-mRNA. Splicing factors are altered in abundance, localization, and activity upon stress, conferring plasticity to the patterns of alternative splicing in response to internal or external stimuli^31–35^. Moreover, chromatin-related factors such as histone modifications and nucleosome positioning, and genomic traits such as gene body length and splice site strength were shown to affect AS30^30, 36–38^.

By employing short-term exposure of Arabidopsis plants to Fe deficiency as a well- explored environmental cue, we report here that the type of gene regulation (i.e., AS or differential expression) is determined by the architecture of the gene, the amplitude of the response, and the temporal pattern of the changes in gene activity. Genomic features are biased towards and typical of specific Fe-responsive processes and seem to govern AS but not transcriptional regulation, which appears to be chiefly driven by physiological requirements. While differential AS (DAS) and differential gene expression (DE) are generally complementary forms of gene regulation, we found this rule suspended when core genes of the Fe deficiency response are considered, suggesting that the two regulatory modes can operate synergistically to allow for an optimally tuned response. We further found that, in many but not all cases, DAS features observed early after the onset of Fe-deficient conditions represent a blueprint of what becomes evident at the activity, transcript, or protein level at later stages of Fe deficiency.

## Results

### Iron deficiency triggers rapid changes in transcription and splicing patterns

To gain insights into transcriptional and post-transcriptional alterations in response to Fe deficiency, Arabidopsis plants were subjected to 0.5, 6, and 12 hours of growth in Fe-free media and subsequent transcriptomic profiling against plants that were transferred to fresh Fe-replete nutrient solution using the RNA-seq methodology. The Illumina HiSeq 4000 sequencing system was adopted to acquire approximately 70-80 million reads for each library with a read length of 100 base pairs (Supplementary Table S1). On average, a total of circa 27,000 genes was expressed in both roots and shoots, of which subsets of 1,161 (roots) and 1,027 (shoots) were defined as differentially expressed between Fe-deficient and Fe-sufficient control plants at one or more of the three sampling time points with relevant expression levels (RPKM > the square root of the mean expression value of the whole dataset), *P* < 0.05, and a fold-change > 2 after normalization using the TMM (Trimmed Mean of M-values) method (Fig. 1a). In both roots and shoots, differentially expressed genes (DEGs) showed a relatively small overlap among the experimental time points, indicative of a highly dynamic and temporally distinctive response to Fe deficiency (Fig. 1c).

**Figure 1.**
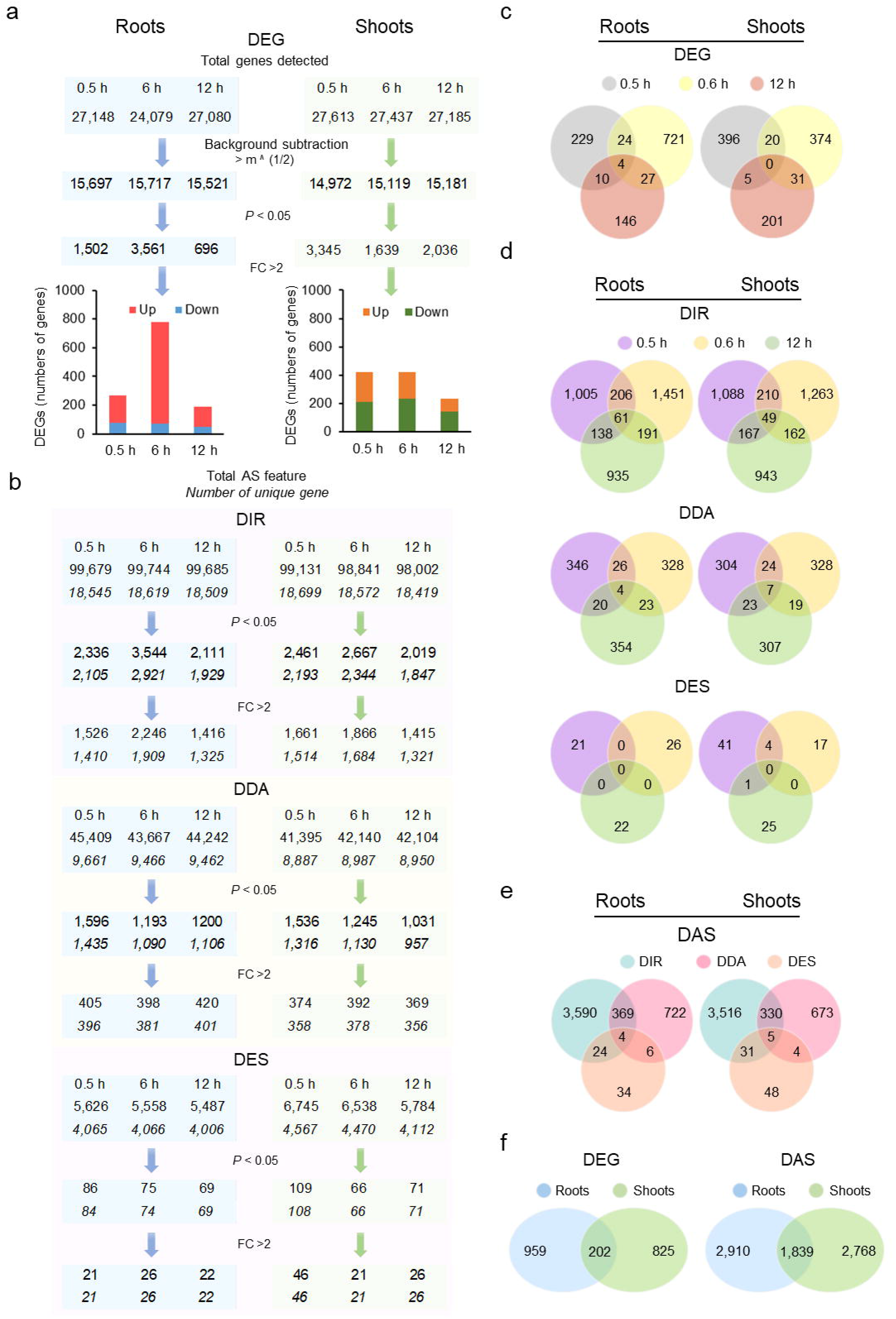
Differentially expressed genes (DEGs) and differentially alternatively spliced (DAS) transcripts in response to short-term exposure to Fe deficiency. a) Filters for the identification and numbers of DEGs at the various experimental time points. b) Venn diagrams showing the overlaps of DEGs among the different time points. c) Filters for the identification and numbers of DAS genes. d) Venn diagrams showing the overlaps of DAS genes among the different time points. e) Venn diagrams showing the overlaps of DIR, DDA, and DES genes in Fe-deficient roots. f) Venn diagrams showing the overlaps of DEG and DAS genes in roots and shoots of Fe-deficient plants. DIR, differential intron retention; DDA, differential donor or acceptor sites; DES, differential exon skipping; FC, foldchange.

Changes in the pattern of alternative splicing (AS) upon Fe deficiency were analysed with the aid of the Read Analysis and Comparison Kit in Java (RACK J) software toolbox^39^. Applying the same thresholds used for the classification of DE genes and considering only genes in which the AS feature was observed in all three replicates, a subset of 7,517 genes was classified as differentially alternatively spliced (DAS), exceeding the number of differentially expressed genes (DEGs) by more than fourfold (Fig. 1f). The largest fraction of DAS genes (82%) produced transcripts that harboured differential intron retention (DIR) features, a smaller subset (15%) carried differential donor/acceptor (DDA) sites, and 2.8% of the DAS transcripts exhibited differential exon skipping (DES) after exposure to Fe deficiency (Fig. 1e). A considerable subset of DAS genes carried two or more different AS features. Genes with DAS features showed distinct overlaps among the different experimental timepoints which depended on the type of AS with DIR being more conserved than DEG and the overlap of DDA genes comparable to that observed for DEGs (Fig. 1c, d). With the exception of DES, in which enhanced and repressed features varied over time, DAS showed a neutral outcome of splicing efficiency upon exposure to Fe deficiency, with about similar proportions of genes carrying enhanced or reduced splicing features (Supplementary Fig. S1).

### Fe deficiency induces a series of concatenated responses in roots and shoots

Transfer of plants to Fe-deplete media triggered the consecutive induction of a suite of distinctly timed responses (Fig. 2). Induction of these processes was almost simultaneously observed in roots and shoots, suggesting that minor changes in Fe supply suffice to sense Fe deficiency in all plant parts. After six hours of exposure to Fe-deficient conditions, induction of the basic Fe uptake machinery comprising the Fe chelate reductase FRO2 and the Fe^2+^ transporter IRT1, vacuolar sequestration of excessive cytosolic levels of secondary-substrate metal cations such as Mn^2+^ and Zn^2+^, and downregulation of Fe import into the vacuole were the most prominent transcriptionally regulated responses in roots (Fig. 2a). Induction of these processes was accompanied by increased expression of a suite of genes encoding regulators such as the *IRONMAN* peptides *IMA1* and *IMA2* and the transcription factor *POPEYE* (*PYE*) (Fig. 2a). A module that was induced slightly later comprised genes mediating the mobilization of Fe in the rhizosphere via secretion of protons and Fe-mobilizing coumarins (Fig. 2b). Induction of Fe mobilisation was accompanied by increased expression of the transcription factors *MYB72* and *MYB10*, which were shown to be crucial for plant survival in soils with severely restricted Fe availability (Fig. 2 b)^40^. Notably, sampling at very early stages of Fe deficiency (0.5 h) revealed expression changes in a direction which was antagonistic to that observed at later stages, suggesting that rearrangements of the transcriptional machinery cause transient perturbations in gene expression that later result in robust transcriptional regulation of these genes (Supplementary Tables S2-5).

**Figure 2.**
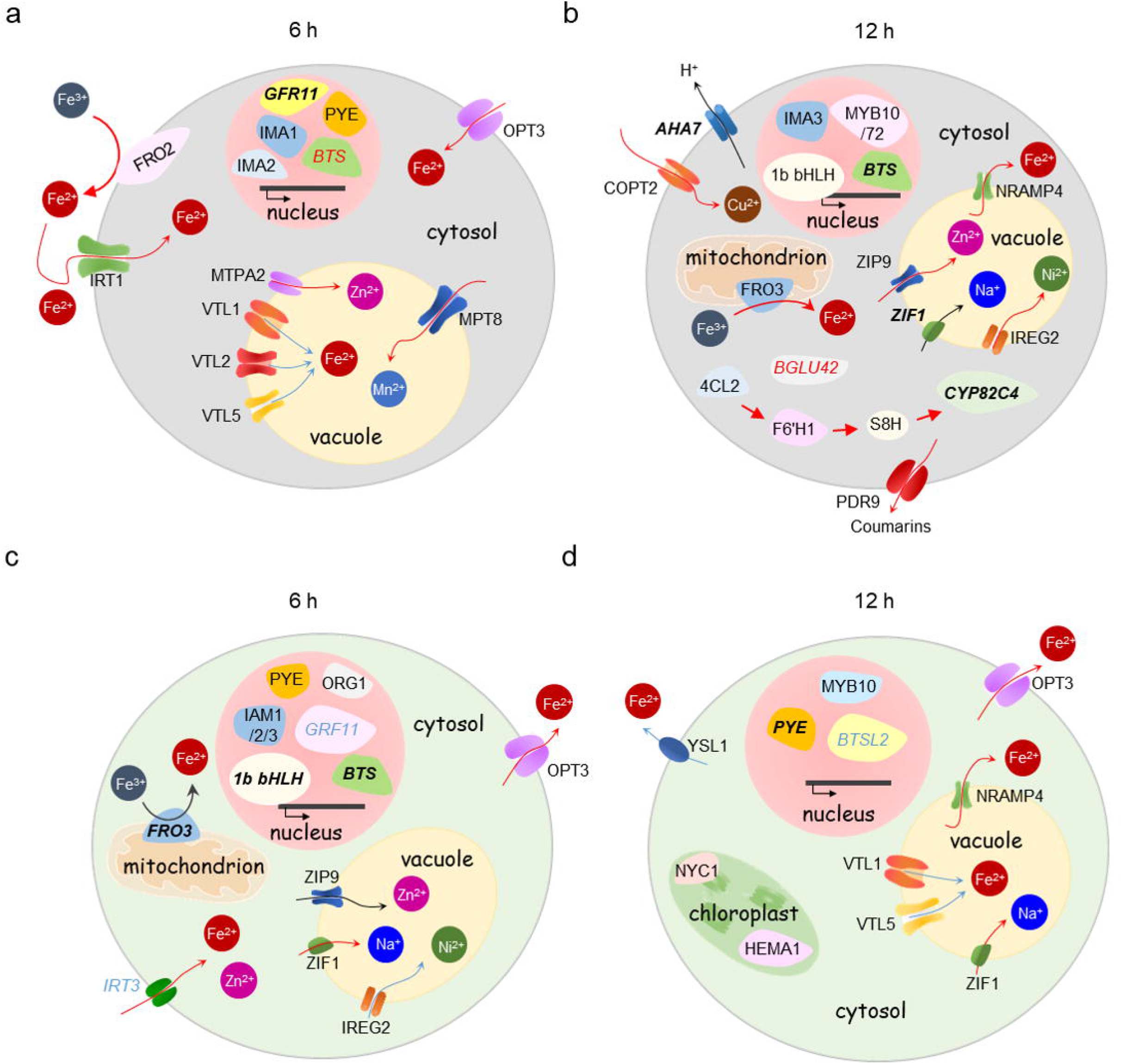
Functions of Fe-responsive genes in Fe uptake and cellular Fe homeostasis. a-d) Cartoon showing DEG and DAS genes in roots (upper panels) and shoots (lower panels) after 6 h (a, c) and 12 h Fe (b, d) deficiency. For the transcriptionally regulated genes, the direction of regulation is indicated by red and blue arrows for up- and down-regulated genes, respectively. DAS genes are denoted by italics, differentially expressed DAS genes are indicated by italics and bold letters. The direction of DAS regulation is indicated by red (enhanced) or blue (reduced) AS. Detailed explanations of gene functions are given in the text.

In shoots, exposure to Fe-deplete media for 6 hours induced Fe transport across the plasma membrane via OPT3 and IRT3 and expression of a variety of regulators such as *PYE*, the E3 ligase *BRUTUS* (*BTS*), subgroup Ib bHLH proteins, and *IMA1*-*IMA3* (Fig. 2c). Similar to root cells, sequestration of transition metals into the vacuole was induced at this time point. In addition, altered transcription of genes involved in ROS homeostasis (*CGLD27*, *ENH1*) was observed in shoots (Supplementary Table S2). After 12 hours, increased trans-plasma membrane transport of Fe-nicotianamine (NA) by YSL1 and Cu^2+^ via COPT2 can be inferred from the induction of the cognate genes at this time-point. Also, chlorophyll metabolism (*HEMA1*, *NYC1*) and ROS homeostasis (*NEET*) were affected by growth on Fe-deplete media (Fig. 2d; Supplementary Table S2).

In addition to genes directly involved in Fe homeostasis, in roots, and to a lesser extent in shoots, short-term exposure to Fe deficiency caused a dramatic increase in the transcription of genes associated with jasmonic acid (JA) biosynthesis and signalling (Fig. 3; Supplementary Table S2 and S3), a response which was strictly restricted to the 6-hour time point. In particular, the first steps of JA biosynthesis and the expression of a suite of transcriptional regulators, referred to as jasmonate ZIM-domain (JAZ) proteins, were strongly induced in response to Fe deficiency (Fig. 3).

**Figure 3.**
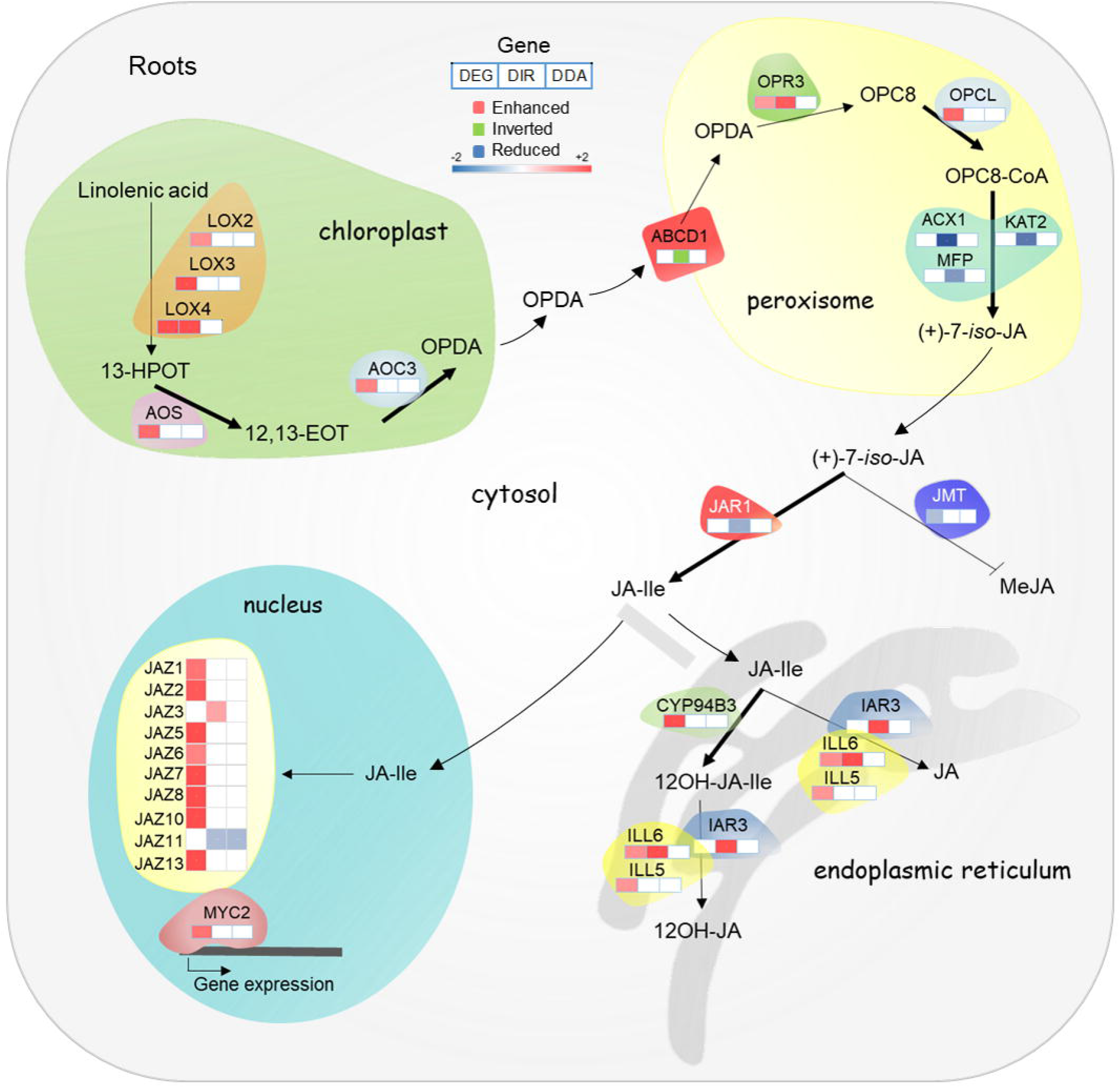
Induction of jasmonate biosynthesis, catabolism, and signalling genes in roots in response to 6 hours Fe deficiency treatment. In plastids, lipid-derived α-linolenic acid is converted by LOX, AOS, and AOC into oxophytodienoic acid (OPDA). Following transport into peroxisomes, OPDA is reduced by OPDA REDUCTASE 3 (OPR3) and converted into JA by β-oxidation. In the cytoplasm, JA is conjugated with amino acids to JA-Ile by JAR1 or to MeJA by JMT. In the endoplasmic reticulum, JA-Ile is degraded to 12OH-JA-Ile and converted to 12-OH-JA via IAR3, ILL5 and ILL6. The latter enzymes can also convert JA-Ile to JA. In the absence of nuclear JA-Ile, expression of JA responsive genes via MYC2 is repressed by JAZ. Upon the entry of JA-Ile into the nucleus, JAZ is degraded via the 26S proteasome and ultimately triggers transcriptional activation of the target genes. Only genes that were differentially expressed or harboured DAS features in response to Fe starvation are shown.

### DAS orchestrates rerouting of the central carbon metabolism

Glycolysis is a key process in energy production and provides metabolic intermediates for biosynthetic processes, storage, and anaplerotic processes. A subset of 75 genes associated with glycolysis was regulated by the Fe regime, almost exclusively by DAS (Supplementary Table S4). Notable exceptions from this pattern were the first and the last step in glycolysis, catalysed by phosphofructokinase (PFK) and pyruvate kinase (PK), which are robustly upregulated at later stages of iron deficiency in transcriptional surveys^9, 11^. Analysis of *PFK1* expression by RT-qPCR showed increasing induction over the first three days of Fe deficiency in roots and, to a much lesser extent, in shoots (Fig. 4a, c). Expression of the Fe status marker *bHLH38* was increased over the first three days of Fe deficiency with a higher transcript level in shoots, indicating that the lower induction of *PFK1* in shoots was not associated with a healthier Fe status of leaf cells (Fig. 4c). Since PFK is catalysing the rate-limiting step in glycolysis, it can be assumed that glycolytic flux is particularly increased in root cells. The final step in glycolysis, the PK-mediated conversion of phosphoenolpyruvate (PEP) to pyruvate, was repressed in roots via enhanced IR of the cytosolic PK isoform At5g08570. In shoots, transcripts of this isoform carried reduced IR upon exposure to Fe deficiency. RT-qPCR analysis revealed decreased transcript abundance of this isoform in both roots and shoots one day after exposure to Fe-deplete media (Fig. 4d). Downregulation of another PK-encoding gene in roots (At3g49160) was reported in several transcriptomic studies^7, 11^, further supporting the supposition that PK activity is decreased upon Fe deficiency.

**Figure 4.**
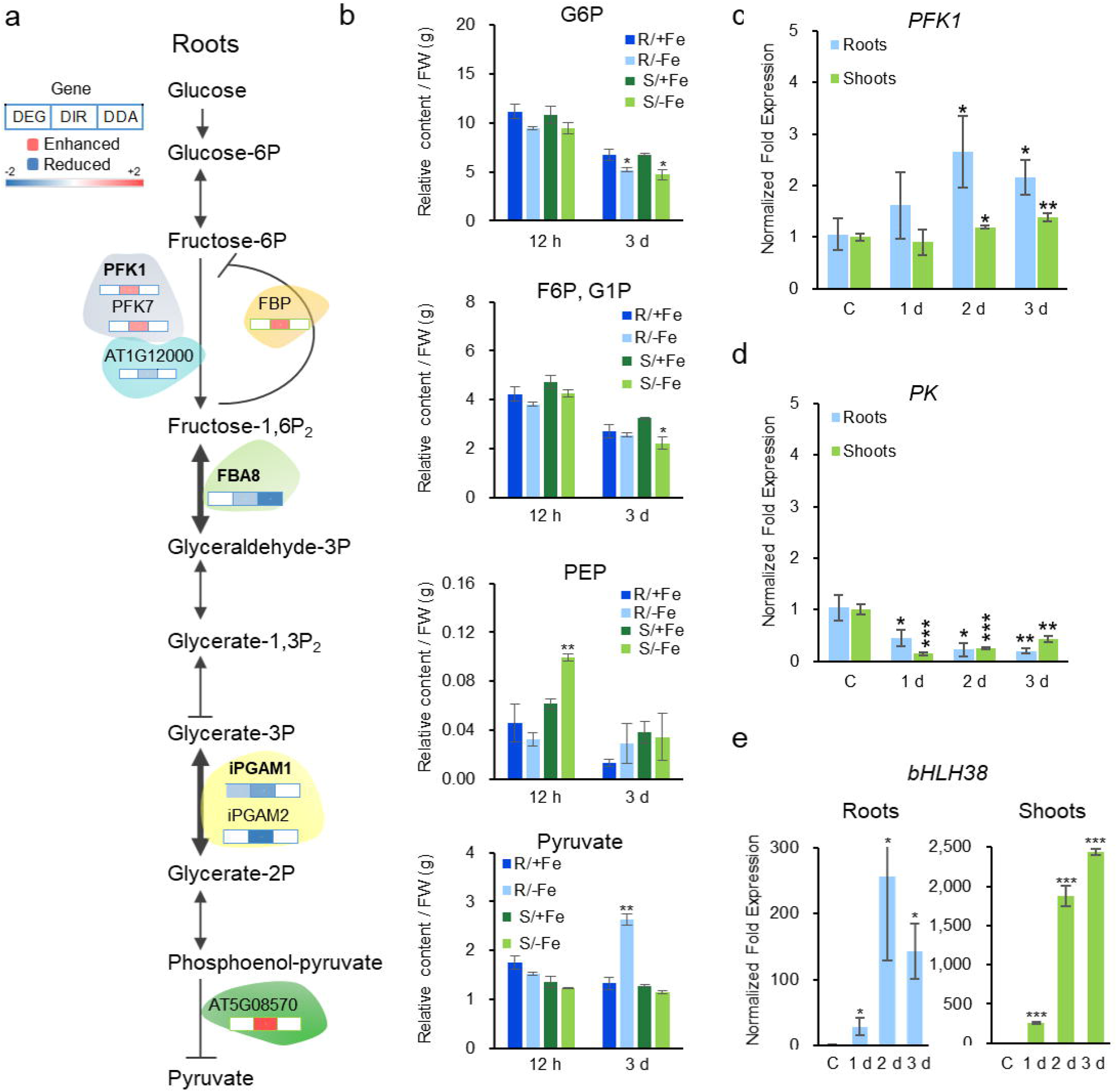
Regulation of glycolytic enzymes and their products by Fe-deficiency. a) DAS regulation of genes encoding glycolytic enzymes in roots. b) Concentrations of glucose-6- phosphate (G6P), fructose-6-phosphate F6P/glucose-1-phosphate (G1P) (G6P and G1P have identical retention times and cannot be distinguished from each other), phosphoenolpyruvate (PEP) and pyruvate in roots and shoots 12 hours and 3 days after transfer to Fe-deplete media. c-e) qRT-PCR analysis of PFK (c), PK (d; At5g08570) and bHLH38 (e). Asterisks indicate significant differences from the wild type in each treatment: *, *P* < 0.05; **, *P* < 0.01; ***, *P* < 0.001. C, control.

For several steps of root glycolysis, DAS appear to precede changes in enzyme activity or abundance. Fructose-bisphosphate aldolase (FBA) catalyses the reversible aldol cleavage of fructose-1,6-bisphosphate, yielding dihydroxyacetone phosphate. In roots, the cytosolic isoform *FBA8* exhibited both repressed IR and DA features (Fig. 4a). Also, the cytosolic phosphoglycerate mutase (PGM) isoforms *iPGAM1* and *iPGAM2* showed reduced IR upon Fe deficiency (Fig. 4a). In a previous study, we observed increased phosphorylation of FBA8 and iPGAM1 protein and accumulation of the (cytosolic) FBA4 protein in roots upon prolonged Fe deficiency^23, 41^, indicating Fe deficiency-induced increase in enzyme activity at later stages of Fe deficiency. Moreover, FBA and PGM accumulated in Fe-deficient roots of *Beta vulgaris* and *Cucumis sativa*^19, 42^, suggesting that increased abundance of these enzymes in response to Fe deficiency is conserved across strategy-I species.

The anti-directional regulation of PFK and PK in roots of Fe-deficient plants implies imbalances of glycolysis intermediates under conditions of Fe starvation. To further investigate this matter, we determined the concentrations of glucose-6-phosphate (G6P), phospho*enol*pyruvate (PEP), and pyruvate over the first three days of Fe deficiency via UHPLC-MS analysis. In roots, but not in shoots, the level of both PEP and pyruvate was increased three days after the onset of Fe deficiency, indicating severe perturbances of pyruvate metabolism at later stages of Fe deficiency (Fig. 4b). Other glycolysis intermediates such as G6P or fructose-6-phosphate (F6P) did neither show pronounced changes in roots nor in shoots upon short-term or extended exposure to Fe deficiency (Fig. 4b), suggesting that the concentrations of theses intermediates are in steady-state during Fe deficiency.

### The TCA cycle malfunctions in roots of Fe-deficient plants

A subset of 34 of TCA-related genes was responsive to the Fe regime (Supplementary Table S5). Similar to what has been observed for glycolysis, genes encoding enzymes of the TCA cycle showed limited transcriptional control and were predominantly regulated by DAS (Supplementary Table S5). In roots, all TCA cycle metabolites under study accumulated three days after the onset of Fe deficiency, with citrate, malate, and succinate being most abundant (Supplementary Fig. S3). In particular, citrate levels were strongly increased; no such accumulation was observed in shoots. Citrate is synthesized in the first committed and pace-making step of the cycle by condensation of oxaloacetate and acetyl CoA, catalysed by citrate synthase (CSY). However, the mitochondrial isoforms *CSY4* and *CSY5* were neither regulated by DAS nor by DE, suggesting other causes for the increased citrate levels.

Aconitase (ACO), catalysing the subsequent conversion of citrate to isocitrate, harbours an active [Fe_4_S_4_]^2+^ cluster, which may compromise ACO activity in Fe-deficient plants and contribute to the accumulation of citrate. In line with this assumption, *ACO2* and *ACO3* were transcriptionally downregulated in roots three days after transfer to Fe-deficient media^7, 9, 43^, indicating restricted conversion of citrate into isocitrate. Also other enzymes of the TCA cycle such as citrate synthase and isocitric dehydrogenase were shown to affect by the iron regime^13^, further supporting the notion that the TCA cycle is compromised or truncated under conditions of Fe deficiency. This scenario is supported by a lack of consistent and mostly repressive DAS-regulation of all steps of the TCA cycle (Supplementary Fig. S3).

### The routes of pyruvate metabolization differ between roots and shoots

Pyruvate derived from glycolysis can be either converted to acetyl-CoA by pyruvate dehydrogenase (PDH) in the mitochondrial matrix, or decarboxylated to acetaldehyde through pyruvate decarboxylase (PDC) in the cytosol. PDH is a multienzyme complex composed of three enzymes, pyruvate dehydrogenase (E1), dihydrolipoyl transacetylase (E2), and dihydrolipoyl dehydrogenase (E3). In roots, isoforms of all three enzymes showed generally reduced IR upon Fe deficiency (Fig. 5a). In shoots, a more complex, mostly repressive regulation was observed (Fig. 5b). Alcohol dehydrogenase (ADH) reduces pyruvate-derived acetaldehyde to ethanol and NAD^+^. In roots, short-term exposure to Fe deficiency resulted in repressed IR of both *PDC1* and *ADH1* transcripts, indicative of increased ethanolic fermentation (Fig. 5a). Also, the class III type alcohol dehydrogenase *ADH2* (*HOT5*) and the putative ADH At4g22110 showed DAS features that were mostly repressed upon Fe starvation. By contrast, in shoots *ADH1* carried chiefly enhanced DIR features (Fig. 5b). Determination of *ADH1* transcript levels in roots by RT-qPCR revealed a steep increase after two days of Fe deficiency (Supplementary Fig. S4), matching transcriptomic studies carried out at later stages on roots of Fe-deficient plants^7, 44^. These data suggest that the ADH-mediated fermentation route is supported in roots, but not in shoots of Fe-deficient plants.

**Figure 5.**
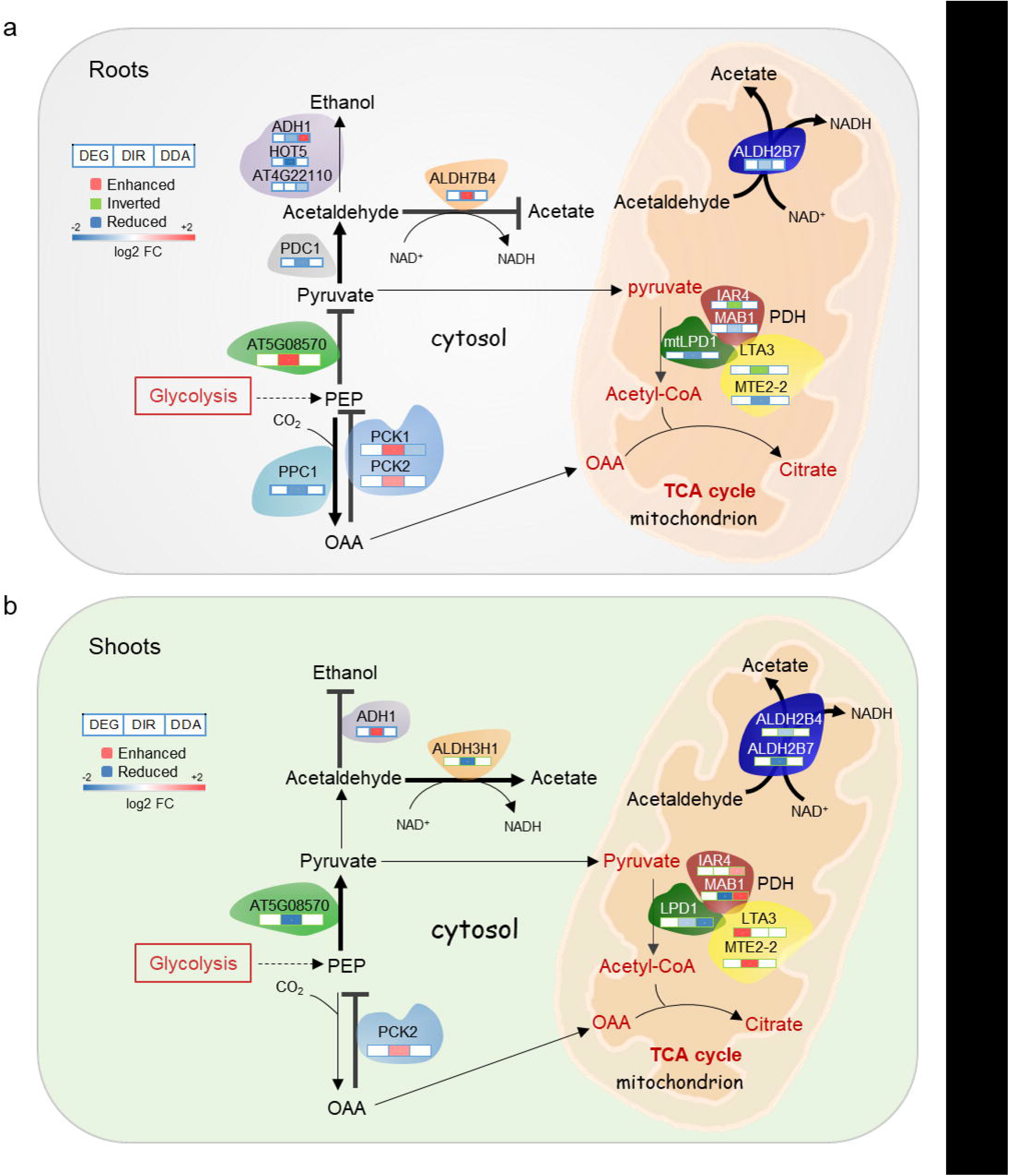
Pyruvate metabolism in roots and shoots of Fe-deficient plants. a, b) Fe-responsive enzymes in roots (a) and shoots (b). Only genes that were differentially expressed or harboured DAS (DIR or DDA) features in response to Fe starvation are shown. See text for details.

An alternative fate of acetealdehyde is the conversion to acetic acid, catalysed by a group of aldehydedehydrogenases (ALDHs) in a reaction yielding NADH. In shoots, three NAD-dependent ALDHs (the mitochondrial family 2 proteins ALDH2B7 and ALDH2B4, and the cytoplasmatic family 3 protein ALDH3H1) showed reduced IR upon Fe starvation (Fig. 5b). In roots, *ALDH2B7* transcripts carried enhanced IR at two time points after the onset of Fe-deficient conditions (Fig. 5a), indicating repressed acetic acid formation. It thus appears that different fermentation routes are engaged in roots and shoots, possibly driven by differences in redox regulation and energy status of leaf and root cells.

Besides reduced PK activity, the build-up of toxic pyruvate concentrations in roots of Fe-deficient plants is circumvented by rapid metabolization of its precursor, PEP. PEP is a potent inhibitor of PFK activity and, if present at high levels, decreases glycolysis rates^45–47^. Increased β-carboxylation of PEP via PEP carboxylase (PPC) in roots is a hallmark of Fe-deficient roots of strategy I plants^17^. In roots, *PPC1* transcripts carried reduced IR features (Fig. 5a), transcription of the gene was induced at later stages of Fe deficiency^7^. The reverse reaction, the conversion of oxaloacetate to PEP by PEP carboxykinase (PCK), showed enhanced IR (Fig. 5a,b). In concert with these results, *PCK1* was found to be downregulated in roots (but not in shoots) of plants subjected to Fe deficiency for three days^7, 48^.

### DE and DAS concertedly regulate the ferrome

Strikingly, the vast majority of DAS-regulated genes did not change significantly in expression over the experimental period. In total, less than 1% of the DAS genes were also differentially expressed (Fig. S1e). Plotting fold-changes of the different DAS types versus DE revealed a moderate correlation between DIR and DE (Fig. 6a). By contrast, regulation of gene activity by DDA and DE was almost mutually exclusive (Fig. 6b).

**Figure 6.**
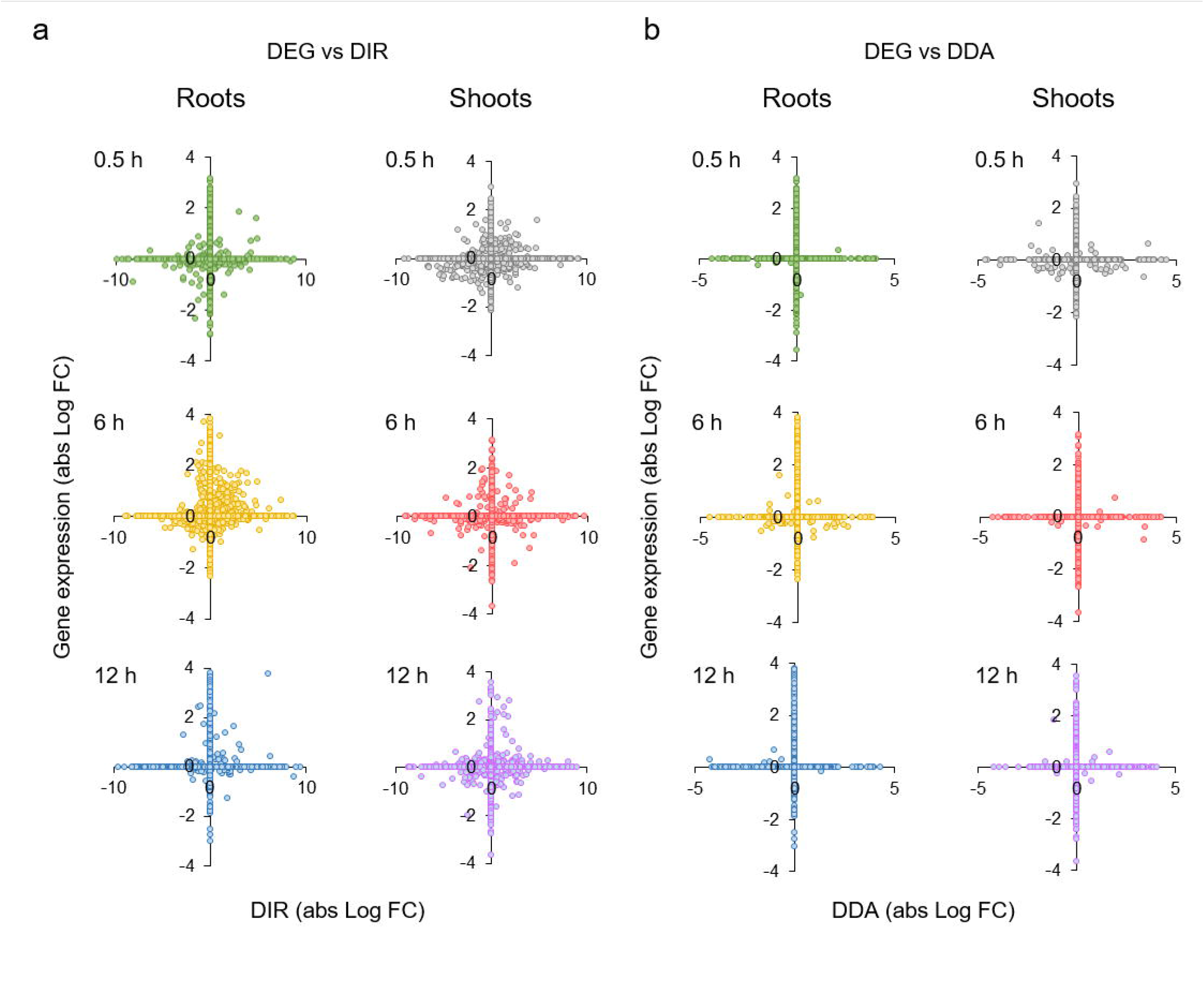
Correlation between DE and DAS of Fe-responsive genes. a) DE versus DIR. b) DE versus DDA.

The principle of mutual exclusivity does not seem to apply when only genes that were shown to robustly respond to Fe deficiency at the transcriptional level in previous studies are considered, a subset that has been referred to as the ‘ferrome’^49^. To revise and update this definition, we surveyed recent public transcriptomic datasets of Fe-deficient plants derived from RNA-seq profiling studies. For roots, seven datasets were analysed, and genes that were found to be differentially expressed in at least four of these studies (three out of five in shoots) were included in the Arabidopsis ferrome. This procedure yielded subsets of 108 and 100 genes in roots and shoots, respectively; a suite of 39 genes was robustly differentially expressed in both roots and shoots (Supplementary Table S6). From 114 DE-regulated ferrome genes, a subsection of 42 (37%) was additionally regulated by DAS (Supplementary Table S2), a percentage that is substantially higher than the average of 3-5% observed in roots at the different sampling points. Most of the genes involved in Fe uptake were transcriptionally regulated; however, some regulators (e.g., *GRF11*, *BSTL2)*, genes encoding enzymes such as the β-glucosidase *BGLU42* which is critical for the deglycosylation of coumarins prior to secretion^50^, and the Zn/Fe transporter *IRT3*, were exclusively regulated by DAS. All of these genes were shown to be transcriptionally induced at later stages of Fe deficiency (Supplementary Table S2).

Unexpectedly, in both roots and shoots the percentage of DE-regulated ferrome genes increased over time. While after 0.5 hours the majority of Fe-responsive genes was DAS- regulated, the proportions of DE and DAS were about equal at 6 hours and massively shifted towards transcriptional regulation at 12 hours (Fig. 7a, b). A comparison with a previous survey of genes responsive to a 3-day-period of Fe starvation revealed a steep decrease in the percentage of DAS-regulated genes at later stages of Fe deficiency (Fig. 7b). Comparing different subgroups within the ferrome showed that transcripts encoding enzymes tended to have a higher percentage of DAS than transporters and regulators. After 3 days, transcriptional regulation of metabolic genes was still lower than in the other categories, but the participation of DAS in the overall gene regulation was dramatically reduced to about relative to what was observed in the short term (Fig. 7b). Such decreased participation of DAS in the regulation of Fe homeostasis was also observed when other gene ontologies were considered (Fig. 7c). This analysis also revealed that at later stages of Fe deficiency transcriptional regulation of ferrome genes is prioritized over other processes, indicating that a more severely disturbed Fe metabolism prioritises expression of a specific subset of genes to secure efficient Fe acquisition. It may also be assumed that at this stage, early DAS events have been established as stable changes in protein abundance, which is not monitored in the two transcriptomic studies considered here.

**Figure 7.**
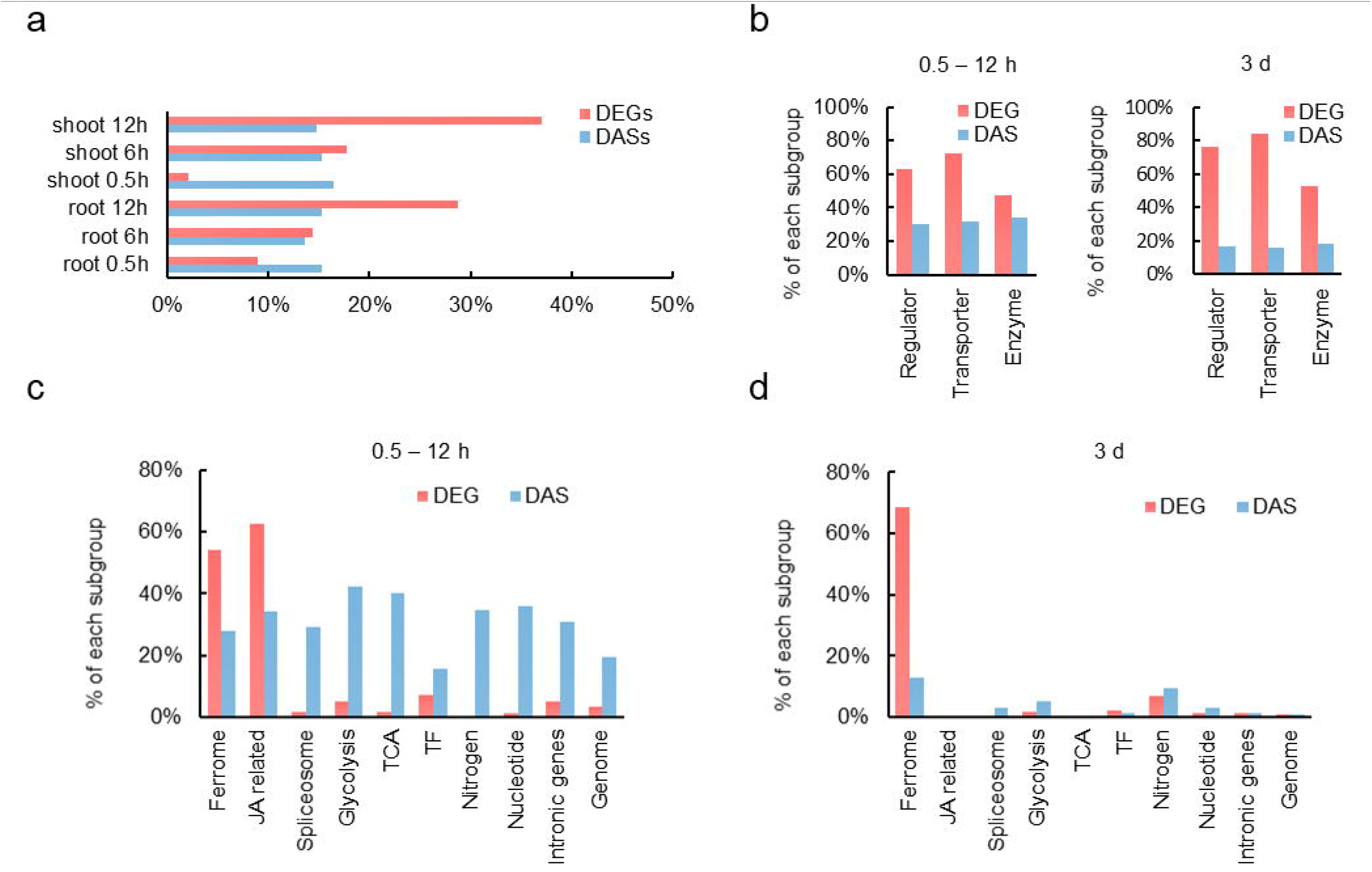
Relative contribution of DE and DAS in the regulation of Fe-responsive genes. a) Time-course of DE- and DAS-regulated genes in roots and shoots in response to Fe deficiency. b) DE and DAS regulation in various categories of ferrome genes after short-tern exposure (left panel) and three days after transfer to Fe-deficient conditions (right panel). Short-term data are pooled from three experimental time points (0.5, 6, and 12 h). Long-term data are taken from a previous study (Li et al., 2014). c) DE and DAS regulation of genes involved in various processes after short-term exposure to Fe-deficient conditions. d) DE and DAS regulation after long-term exposure (3 d) to Fe-deficient conditions.

### Gene architecture predefines the type of gene regulation

From the survey of Fe-responsive genes, it appears that the various Fe-responsive processes are preferentially controlled by either DE (e.g., JA-biosynthesis and signalling), DAS (glycolysis and TCA cycle), or both DE and DAS (Fe uptake), regulatory patterns that are possibly associated with the amplitude of the response (Supplementary Tables S2-5). To investigate whether genomic traits influence or govern the type of gene regulation, we first analysed the number and length of introns of Fe-responsive genes. Intron length was only moderately correlated with the probability of IR. Introns with a length of more than 200 bp had a slightly higher chance of being retained upon Fe deficiency than shorter introns, reaching a peak at about 700 bp (Supplementary Fig. S2a). The number of introns, on the other hand, appeared to be a strong predictor for AS. When normalized to the average intron number across the genome, the number of genes producing transcripts with DIR, DDA, or DES features showed a steep incline which saturated at an intron number between 10 and 20 (Supplementary Fig. S2b, c). Notably, intron-rich genes harbouring >50 introns were rarely found to be subject to DAS (Supplementary Fig. S2b, c).

As further parameters that presumptively affect the type of gene regulation, we investigated the influence of 5’ and 3’ splice site strength, promoter length, and the number of transcription factor binding sites (TFBSs) on DE and DAS. Plotting the minimum average splice site strength [ASS; (min 5’ + min 3’)/2] of Fe-responsive genes against the intron number showed that weak splice sites were generally associated with lower intron numbers (Fig. 8a-c), a finding that may be causal rather than merely correlative. Considering DE- and DAS-regulated genes separately revealed that the average intron number was much lower in DE than in DAS genes, while ASS appears to be biased towards higher values (i.e., stronger splice sites) in this group relative to DAS-regulated genes. With few exceptions, genes with intron numbers >20 were regulated by DAS (Fig. 8b). However, highly negative ASS values do not appear to compromise transcriptional regulation, indicating that DE can be promiscuously employed to regulate genes hardwired for being DAS-regulated. Factors that may support transcriptional regulation such as promoter length and the number of transcription factor binding sites (TFBSs) did not pose major effects on gene expression.

**Figure 8.**
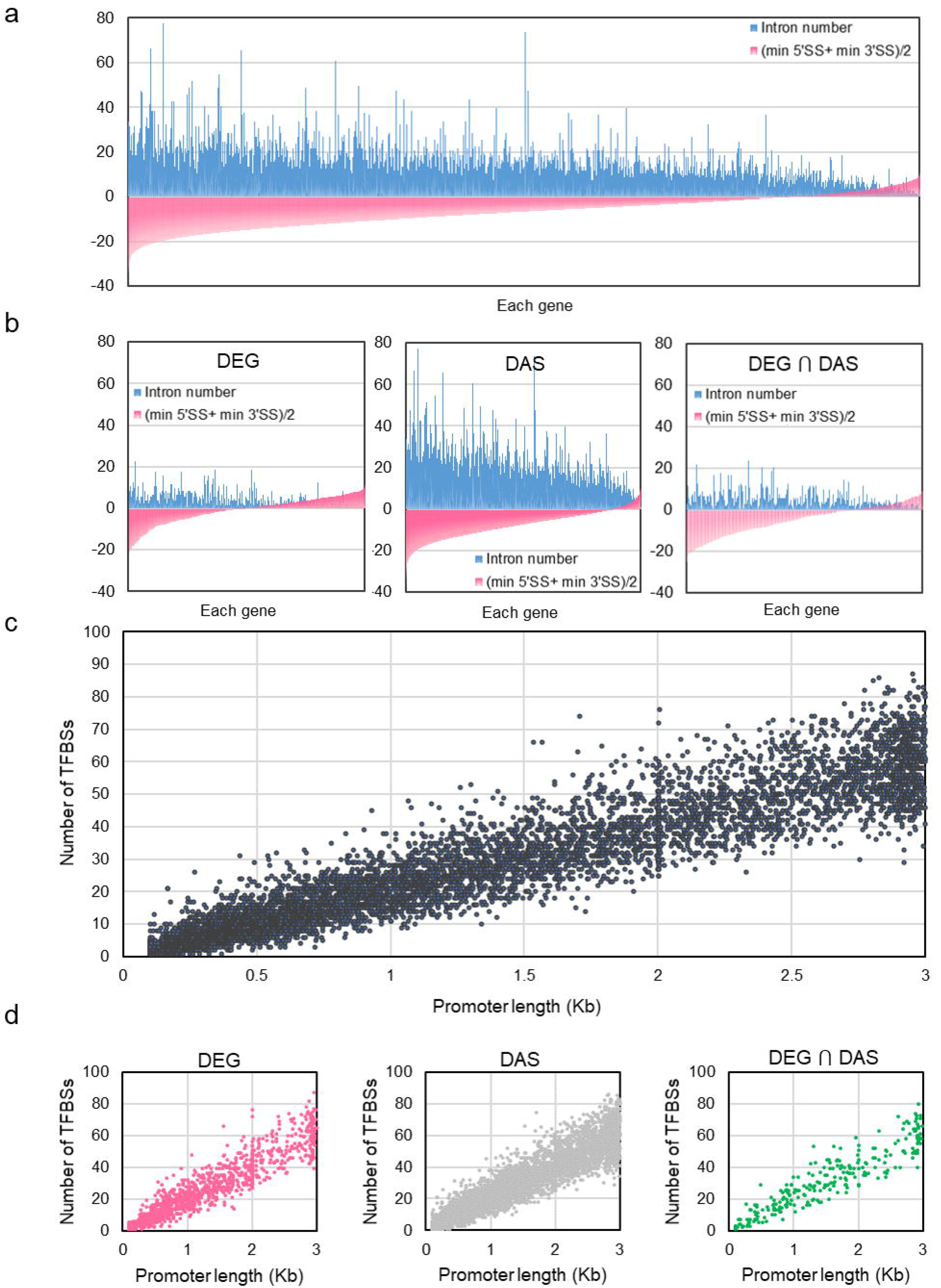
Correlation of architectural traits of Fe-responsive genes with the mode of gene regulation. a, b) Intron number (blue bars) and average minimum 5’ and 3’ splice site (SS) strength (red bars) of all Fe-responsive genes (a), and DEGs, DAS genes, and genes that are regulated by both DE and DAS (b). c, d) Correlation between the number of transcription factor binding sites (TFBSs) and promotor length of all Fe-responsive genes (a), and DEGs, DAS genes, and genes that are regulated by both DE and DAS (b).

While promotor length was correlated with the number of TFBSs, no bias towards higher values was observed for DE-regulated genes, suggesting that these parameters are not decisive for the type of regulation (Fig. 8c, d). It thus appears that DE is not dictated by genomic features and can be called into play when more robust regulation is required.

### The type of gene regulation is genomically hardwired across biological process

To further investigate the mechanisms underlying the regulation of Fe-responsive genes, we compared the genetic architecture of genes involved in various processes that are affected by the Fe regime. Homing in on individual genes of the ferrome, a comparison of the DE- and DAS-regulated subsets supported the trend observed for all Fe-responsive genes, i.e., a marked shift towards stronger splice sites for DE-regulated genes (Fig. 9). However, regulation by DE was also observed for genes with very weak (i.e., highly negative) splice sites and high intron numbers (Fig. 9a). The difference in ASS between DE- and DAS-regulated genes was obvious at all experimental time points, and was independent of the plant part under study (i.e., roots vs. shoots) and the direction of regulation (Supplementary Fig. S3).

**Figure 9.**
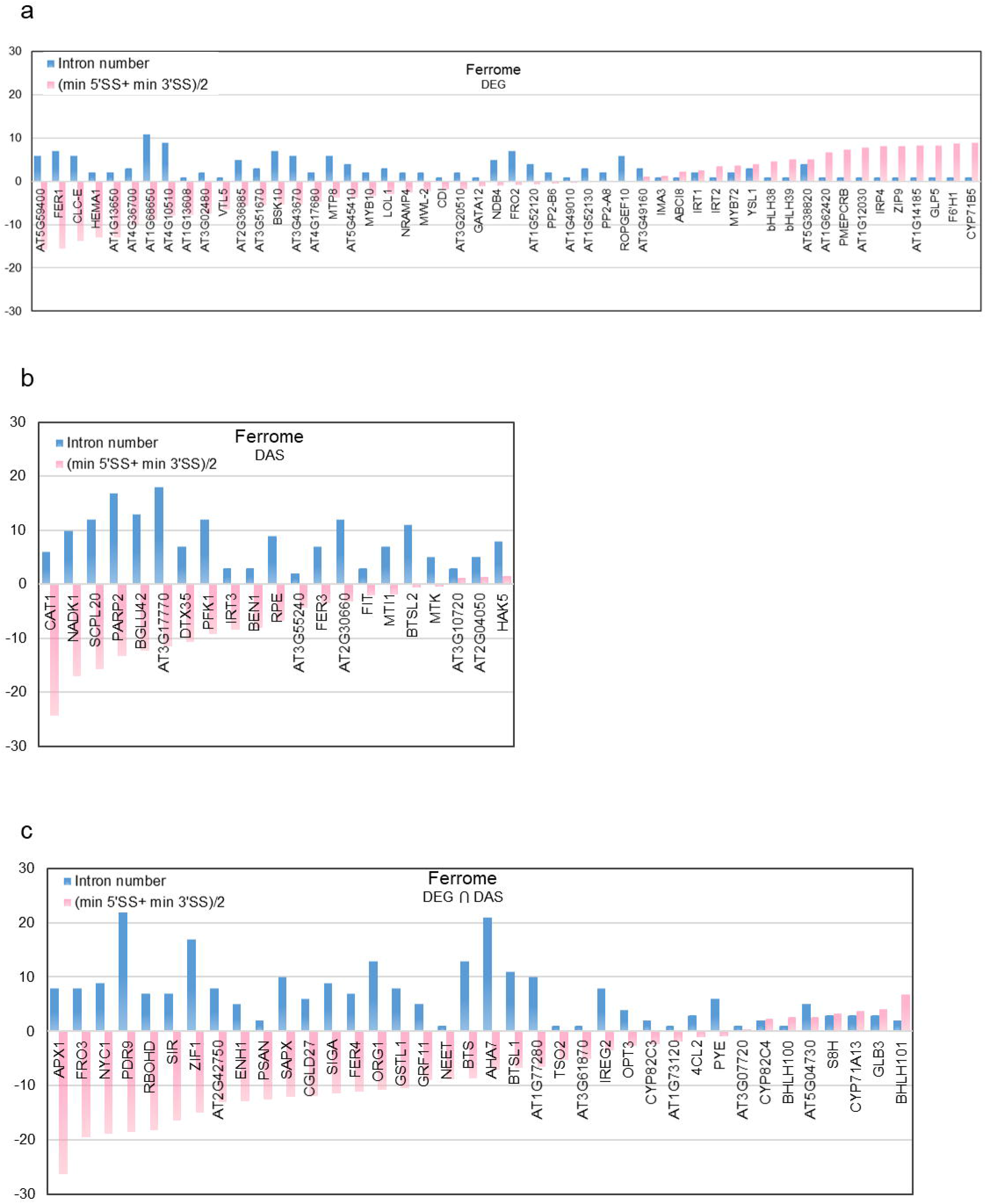
Correlation of architectural traits of individual ferrome gene with the mode of gene regulation. a-c) Intron number (blue bars) and average minimum 5’ and 3’ splice site (SS) strength (red bars) of DEGs (a) DAS genes (b), and genes that are regulated by both DE and DAS (c).

In addition to the ferrome, Fe-responsive genes involved in glycolysis (75 out of a total of 133), the TCA cycle (34/63), JA-signalling (25/36), transcriptional regulation (432/1717), and pre-mRNA splicing (163/396) were analysed. This comparison revealed pronounced differences in splice site strength and intron number among the various groups that appear to be typical of a particular process. When compared with all Fe-responsive entities (*n* = 9,190), genes from the ferrome group, JA-related genes, and genes encoding transcription factors showed a lower-than-average number of introns and a higher-than-average ASS, while genes related to glycolysis, the TCA cycle, and splicing-related genes exhibited an opposite pattern (Fig. 10).

**Figure 10.**
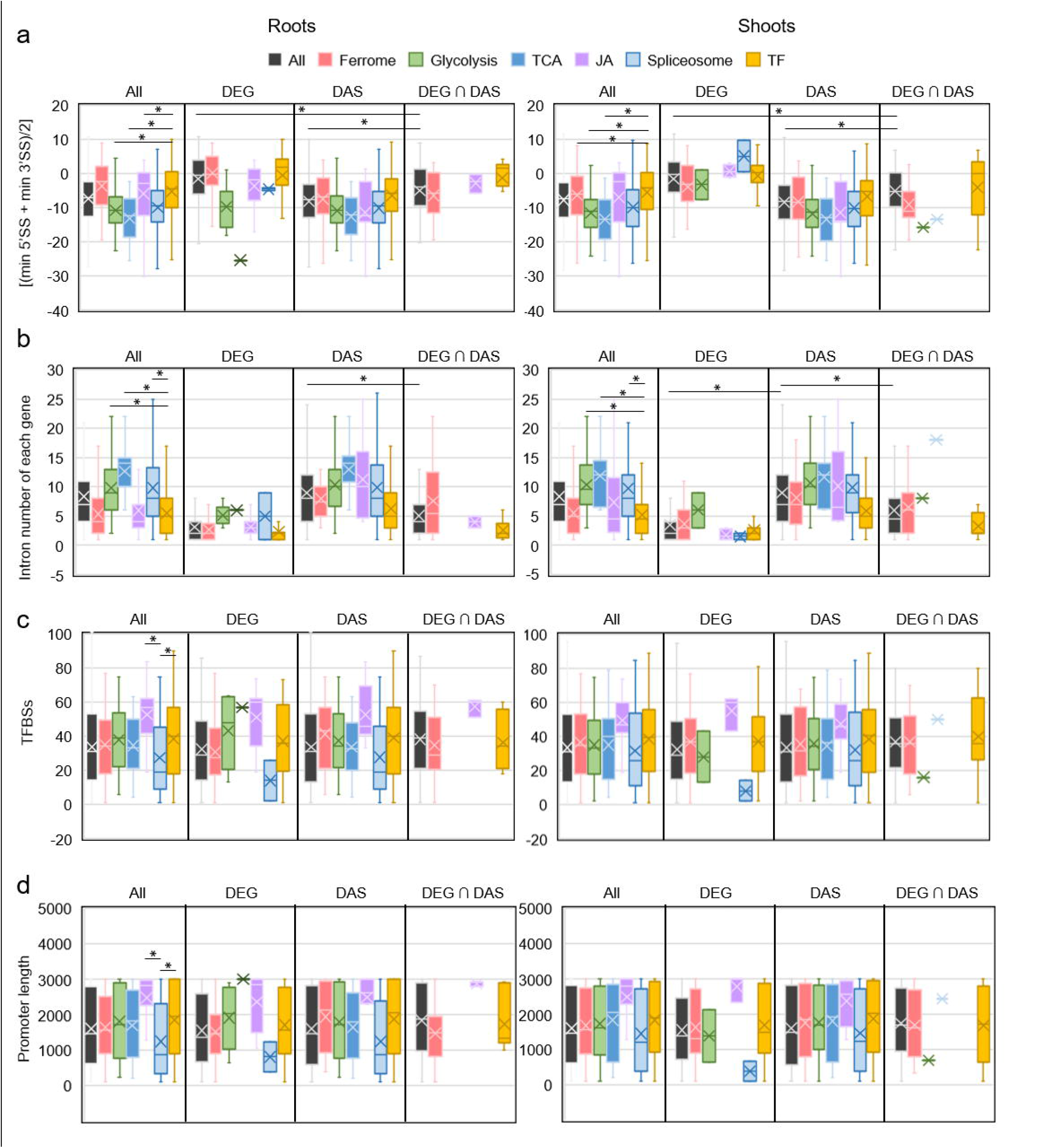
Genomic features of Fe-responsive genes involved in various processes. a) Average minimum strengths of 5’ and 3’ splice sites. b) Intron number. c) Transcription factor binding sites (TFBSs). d) Promoter length. Box plot shows the median (line) and the average (x) for genes in roots (a) and shoots (b). Significant differences were detected using two-way ANOVA with Tukey’s multiple comparison test, *P* < 0.001.

Separating exclusively DE- or DAS-regulated genes revealed pronounced differences between the two groups, with DE-regulated genes exhibiting much stronger splice sites and lower intron numbers (Fig. 8b). Strikingly, DAS-regulated transcription factors harbour features of DE-controlled genes, possibly reflecting an evolutionary trend of this group towards this type of regulation. It is further interesting to note that – while containing transcription factors and metabolic genes to almost equal portions – ferrome genes behaved similar to genes encoding transcription factors, suggesting that the massive changes in abundance in response to environmental cues necessitates this type of regulation in most genes in these two groups to allow for an adequate and efficient response to environmental cues.

Although promoter length and the number of TFBSs do not seem to affect the mode of gene regulation, these parameters can be supportive to or even critical for gene expression. For example, in yeast, promoters of stress-responsive genes were found to be longer than those of housekeeping genes^51^, suggesting that such responsiveness requires a more elaborate interaction between *trans*-acting factors and *cis*-regulatory elements. In the present study, the average promoter length (1,595 bp) and the average number of TFBSs (33.5) of all 9,190 Fe-responsive genes were not significantly different from those determined for genes of the categories ferrome, glycolysis, and TCA cycle, but slightly higher (38.1) for genes encoding transcription factors (Fig. 9). Crucially, splicing-related genes had on average significantly shorter promotors (1,290 bp) and less TFBSs (27.5), indicating a trend against regulation by DE of these genes. Another deviation from the overall average was observed for JA-related genes, which had longer promoters and more TFBSs (Fig. 11). The promoter architecture of JA-related genes may reflect the extent of recruitment by different signalling pathways. For example, genes encoding enzymes mediating the first steps of JA synthesis (i.e., *LOX2*, *LOX3*, *LOX4*, *AOS*, and *AOC3*) had an average promotor length of 2,764 bp and 63.6 TFBSs, almost twice of the average of Fe-responsive genes. JA synthesis is important for a plenitude of processes, which is possibly mirrored by a large number of regulatory *cis*-consensus motifs on the promotors of these genes. By contrast, clade Ib bHLH proteins, which are highly responsive to the Fe regime but have not been associated with other responses, have short promotors (on average 918 bp) and relatively few TFBSs (24.5). Collectively, the data suggest that a particular process is prone to a certain type of gene regulation, which is, by an appreciable extent, hardwired by the structure of the genes involved in this process.

**Figure 11.**
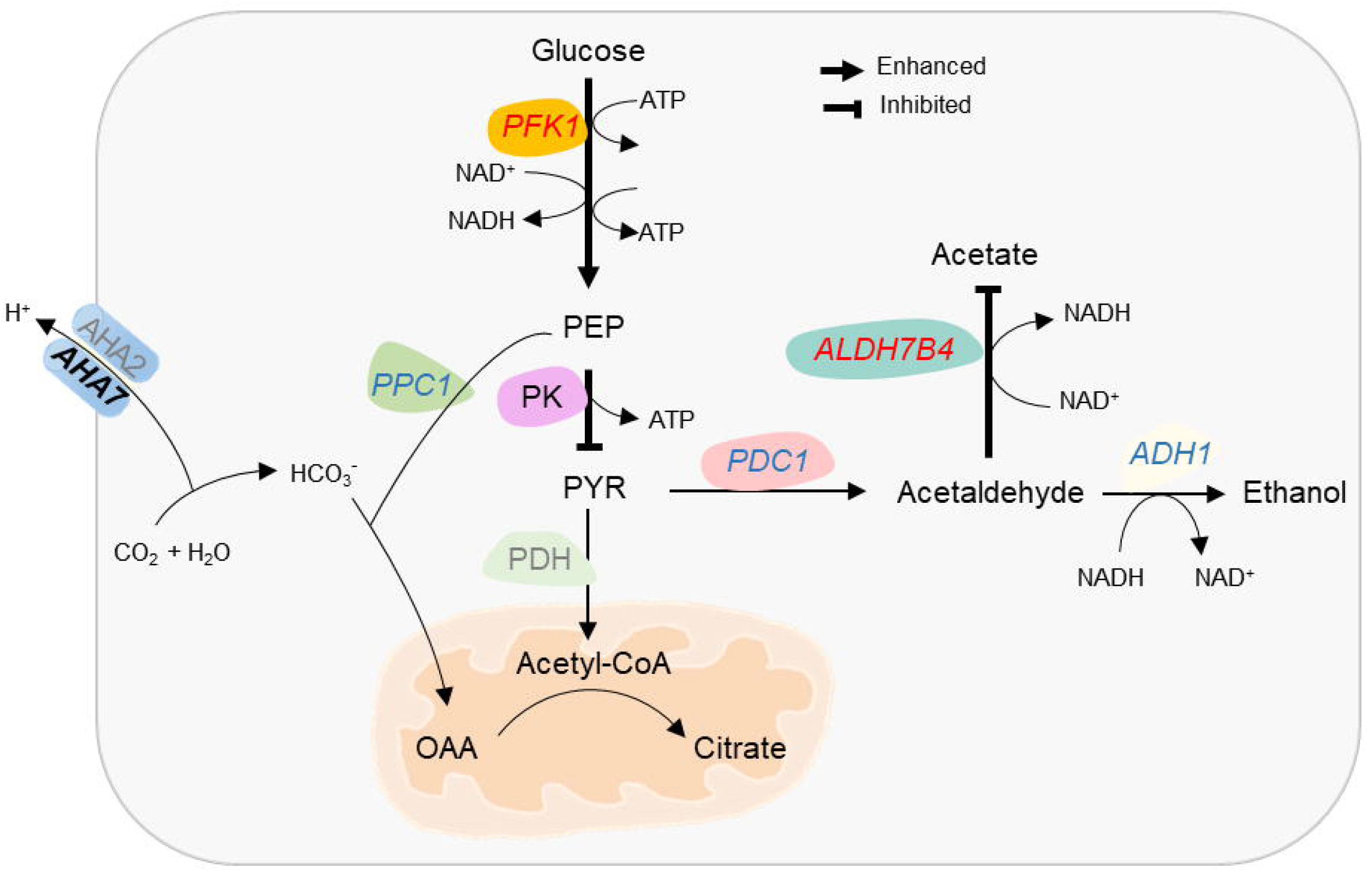
Summary of the changes in central carbon metabolism in roots of Fe deficient plants. Carbon flux through the glycolytic pathway is increased by transcriptional upregulation of *PFK1*. Pyruvate accumulation caused by a truncated TCA cycle is avoided by downregulation of PK activity by DAS and DE and increased ethanolic respiration via PDC1 and ADH1, which are regulated by DAS and DE. This reaction regenerates the NAD^+^ pool for continued glycolytic flux. Acetate respiration, which is prioritized in shoots, is repressed by DAS of *ALDH7B4*. Increased net proton secretion by P-type ATPases is counteracted by the formation of HCO_3_ and subsequent carboxylation of phospho*enol*pyruvate (PEP), yielding oxaloacetate (OAA), which is converted to citrate. DAS genes are denoted by italics, genes regulated by both DE and DAS are indicated by italics and bold letters. Enzymes that are not differentially expressed are depicted in grey. The direction of DAS regulation is indicated in red (enhanced) or blue (reduced) letters.

## Discussion

### Multi-layered gene control orchestrates the acclimation to Fe deficiency

The current survey shows that short-term exposure to Fe deficiency is sufficient to trigger profound changes in mRNA abundance and AS patterns, inducing a series of responses that act in concert to recalibrate cellular Fe homeostasis. These responses comprise alterations in transport processes, secondary metabolism, redox and pH balance, hormone signalling, and central carbon metabolism, and are simultaneously induced in roots and shoots. A less well-explored component Fe deficiency response is the sharply timed induction of JA biosynthesis. Similar to what we describe here for the strategy I plants Arabidopsis, short-term activation of JA biosynthesis was reported for the strategy II plant rice^52^, suggesting that expression of JA-related genes constitutes a general, conserved mechanism across land plants. The reasons for this transient upregulation of JA synthesis are yet to be elucidated. Activation of JA signalling by exogenously supplied methyl JA was shown to affect the assembly of the microbial community in the rhizosphere, possibly by altering the composition of root exudates^53^. In addition, JA determines the compatibility between host and beneficial microbes^54,55^, which, in turn, may positively affect coumarin-mediated Fe uptake^56^. An alternative explanation can be inferred from a study on Arabidopsis plants exposed to oxygen deficiency. Here, a similar boost in JA production was observed in roots after a 6-hour period of oxygen depletion and suggested to trigger repression of root meristem regulators to avoid energy exhaustion when respiration is compromised^57^, a scenario which may also apply to Fe-deficient plants.

An underappreciated response to Fe deficiency is the reprogramming of large parts of the central carbon metabolism to counteract imbalances resulting from reduced respiration, decreased activity of Fe-containing enzymes, and shifts in intracellular redox and pH homeostasis. Rerouting central carbon metabolism to prioritize Fe uptake under conditions of Fe deficiency was observed in soil bacteria^58^ and eukaryotic systems such as human macrophages^59^ and yeast^15, 60, 61^, suggesting that alterations in primary metabolism represent a conserved concept across organisms. Notably, the priority for anaerobic glycolysis, inhibition of PK activity, and the truncated TCA cycle observed in Fe-deficient plant cell resembles the metabolism of cancer cells^62^, further underlining the plasticity of central carbon metabolism and its role in the adaptation to a given set of conditions. Since many of the observed perturbations such as, for example, the accumulation of pyruvate or citrate are undetectable or at least not pronounced within the first 12 hours of Fe deficiency, it can be assumed that alterations in gene activity occur both in response to and in anticipation of imbalances caused by discontinued Fe supply.

### Pyruvate is a central player in the metabolic homeostasis of Fe-deficient plants

High pyruvate levels stimulate respiration^63^, a situation which is not desirable under conditions of Fe-deficiency. In roots of Fe-deficient plants, prevention of such build-up is attempted by inhibition of PK and induction of ADH. ADH transcript and protein levels were found to increase in Fe-deficient roots of various species^7, 23, 42, 44, 64, 65^, suggesting that ethanol fermentation is a common feature of the Fe deficiency response. In shoots, however, acetic acid is a more likely product of anaerobic respiration. The different fate of pyruvate in root and leaf cells may be associated with different redox status of root and leaf cells. In roots, reduction of acetaldehyde to ethanol restores the NAD^+^ pool to maintain high glycolysis rates^66^. In Fe-deficient leaf cells, acetate synthesis may recalibrate redox homeostasis, a scenario which was suggested to constitute a critical component of a survival strategy triggered by drought stress and possibly other environmental changes^67–69^. In this strategy, acetate formation via ALDH2B7 is crucial in counteracting oxidative stress by stimulating COI1-dependent jasmonate signalling and subsequent induction of JA-responsive genes^69^.

Another peculiarity associated with the metabolism of Fe-deficient root cells is related to the massively increased export of protons, which necessitates strategies to replenish substrate for the H^+^-ATPases and to prevent excessive alkalisation of the cytosol. The concentration of free cytosolic H^+^ is in the sub-micromolar range^70^ and insufficient to support the high proton fluxes of Fe-deficient root cells. Such compensatory release of protons is achieved by PPC-mediated carboxylation of PEP or, more precisely, the preceding formation of HCO_3_^-^ by carbonic anhydrase, which is accompanied by a net production of protons (Fig. 11). The product of PEP carboxylation, OAA, was not detectable in roots, indicative of its rapid metabolization to citrate, the level of which was dramatically increased upon Fe starvation. Citrate accumulation in roots of Fe-deficient has been reported for a variety of species with high proton extrusion capacity such as tomato^16^, cucumber^71, 72^, sugar beet^73^, and *Capsicum annuum*^74^, implying a link between these two observations. Moreover, citrate levels appear to correlate with PPC activity^75^, making it tempting to speculate that increased proton secretion and subsequent PEP carboxylation are the driving forces for citrate accumulation in Fe-deficient roots. In Arabidopsis, PPC1 was found to be induced in proteomic^43^ and transcriptomic surveys of Fe-deficient roots^7, 9, 43, 76, 77^. Moreover, transcriptional induction of PEP carboxylase kinase (*PPCK1*)^7, 9, 77, 78^, as well as phosphorylation of PPC1 was observed in Fe-deficient roots^23^, indicating multi-layered and robust regulation of PEP carboxylation in roots upon Fe starvation. It can thus be assumed that citrate accumulation is caused by increased production of OAA via PEP carboxylation and limited metabolization of citrate due to compromised activity of the Fe-sulphur proteins aconitase and succinate dehydrogenase, compromising the completion of the TCA cycle. In Arabidopsis, ATPase-mediated proton export is regulated by FIT^79^, which would imply that citrate accumulation is also dependent on functional FIT protein. In line with this concept, no increased citrate levels upon Fe deficiency were observed in *fit* mutants^80^. Taken together, these data imply that, with a few exceptions where such changes are induced at the level of transcription (e.g., *PFK*, *PK, ADH,* and *PPC*), DAS appears to represent the main regulon of central carbon metabolism, in particular at early stages of Fe deficiency. This pattern differs from that observed for genes encoding enzymes involved in the production of Fe-mobilizing coumarins, which massively change in abundance both at the transcript and protein levels (this study)^23, 81^. Thus, metabolic processes are not necessarily associated with DAS regulation; it rather appears that the amplitude and specificity of the response (i.e., the direct involvement in Fe uptake) determine the mode of gene regulation.

### Transcription is biased by but not dependent on genomic features

A recent analysis of a suite of AS datasets derived from various eukaryote systems revealed that in addition to or in conjunction with splicing factors, genomic traits may determine the propensity of the mode by which a gene is regulated, suggesting that such features hardwire genes to be controlled by either transcription or alternative splicing^30^. In support of this conception, we show here that in particular the number of introns and the strength of the splicing sites influence the probability of a gene to undergo AS. In contrast to DAS, transcriptional regulation does not appear to be dependent on genomic features. While exclusively DE-regulated genes are strongly biased towards low intron numbers and high splice site strengths, these traits are rather compromising DAS regulation than representing a prerequisite for transcriptional regulation. Transcription appears to be promiscuously employed when pronounced changes in gene activity are required. A representative example is *PFK1*, which, despite having the architecture typical of DAS-regulated genes and being part of a chiefly DAS-regulated process, is robustly controlled by transcription. It should be noted, however, that transcription-promoting features such as the number of TFBSs may bias the mode of gene regulation towards, or, as in the case of splicing factors, against regulation by DE. The generally observed mutual exclusivity of DE and DAS was not evident in ferrome genes, suggesting that the rapid and highly dynamic responses to the availability of Fe requires a more complex regulation to avoid extreme depletion of or sudden overload with Fe. Thus, under a certain set of conditions, the two mechanisms appear to collaborate to regulate gene activity. Mostly, but not always, such cooperative regulation occurs anti-directionally, suggesting that DAS is fine-tuning translation efficiency.

## Conclusions

Collectively, our data show that DE and DAS are co-operative mechanisms that jointly, but mostly autonomously, govern gene activity in a precisely timed pattern in response to environmental signals. The current survey further revealed components of the Fe deficiency response that are either not evident under steady-state conditions (i.e., JA signalling), or not sufficiently mirrored at the transcript level and thus underappreciated in transcriptomic studies (i.e., alterations in pyruvate metabolism). Our data further show that DAS events triggered by short-term exposure to Fe starvation are in large part congruent with observations derived from proteomic, physiological, and metabolic studies conducted at later stages of Fe deficiency, forerunning subsequent alterations in protein abundance and enzyme activity. AS appears to represent an area of largely unexplored ‘dark matter’, controlling putatively important responses that may significantly contribute to the pronounced discordance of mRNA and protein expression, a gap that is particularly wide in plants^82^. Thus, DAS can be considered as a major contributor of such discordance and a putative proxy for more robust metabolic or physiological changes. A surprising finding was the strong bias towards the mode of gene regulation posed by genomic features. It seems reasonable to speculate that particular processes are more efficiently regulated by either regulatory mode, depending on the function of the genes and the amplitude of the alterations in gene activity required for adequate acclimation to adverse environmental conditions.

## Materials and Methods

### Plant Growth

Seeds of Arabidopsis (*Arabidopsis thaliana* (L.) Heynh) accession Columbia (Col-0) were obtained from the Arabidopsis Biological Resource Center (Ohio State University). Plants were grown hydroponically in a nutrient solution composed of 5 mM KNO_3_, 2 mM Ca(NO_3_)_2_, 2 mM MgSO_4_, 2.5 mM KH_2_PO_4_, 14 μM MnCl_2_, 70 μM H_3_BO_3_, 1 μM ZnSO_4_, 0.5 μM CuSO_4_, 0.2 μM Na_2_MoO_4_, 0.01 μM CoCl_2_, 40 μM Fe-EDTA, and 4.7 mM MES buffer (pH 5.5). Seeds were infiltrated with distilled H_2_O for 3 days in dark in 4°C before being transferred to the hydroponic system and then grown in a growth chamber at 21°C under continuous illumination (7 70 μmol m^−2^ s^−1^). After 16 d of pre-cultivation, plants were transferred to fresh nutrient solution with either 40 μm Fe-EDTA (+Fe plants for control) or no Fe with 100 μM 3-(2-pyridyl)-5,6-diphenyl-1,2,4-triazine sulfonate (-Fe plants) for 0.5-, 6-, and 12-hours treatment for RNA-seq analysis, and 1-3 days -Fe treatment for further RT-qPCR experiments or Ultra-High Performance Liquid Chromatography (UHPLC) analysis. At the end of the treatment, root and shoot were collected and stored at -80°C.

### RNA-seq and definition of DEGs

Total RNA was extracted from approximately 100 mg of Arabidopsis roots or shoots using the RNeasy Plant Mini Kit (Qiagen, Cat. No. 74904). RNA samples were treated with DNaseI (Qiagen, Cat. No. 79254) to remove DNA. RNA concentration was determined with a NanoDrop ND-1000 UV-Vis spectrophotometer (NanoDrop Technologies). For preparing RNA-seq libraries, mRNA molecules with poly-A tails were purified using poly-T oligo-attached magnetic beads.

The first-strand cDNA was synthesized by the use of dNTP (dUTP replaced by dTTP), buffer, RNaseH, and DNA polymerase I. cDNA was purified using a Purification Kit (Qiagen) followed by performing end repair and A-tailing. The sample was then treated with USERTM (Uracil-Specific Excision Reagent) enzyme to digest the antisense strand DNA follow by PCR reaction. After these procedures, the library could be sequenced using the Illumina HiSeq 4000 platform. The first step in the trim process was the conversion of the quality score (Q) to error probability. Next, for every base a new value was calculated; 0.05 – error probability. This value is negative for low quality bases, where the error probability is high. For every base, we calculated the running sum of this value. If the sum dropped below zero, it was set to zero. The part of the sequence to be retained is between the first positive value of the running sum and the highest value of the running sum. Everything before and after this region was trimmed off. In addition, reads shorter than 35 bp were discarded. A total of 66 to 87 million reads were obtained from Illumina sequencing for the various libraries (Supplementary Table S1). Reads were aligned to the TAIR10 transcriptome using Bowtie2^83^, and only alignments of read pairs that mapped to the same transcripts were accepted. The remaining reads were mapped to the TAIR10 genome directly using the BLAT program^84^ with default parameters. Alignments with a minimum 95% identity for each read were considered for mapping but only the alignment with the highest identity were accepted. Read counts were computed using the RACKJ software package (http://rackj.sourceforge.net/), normalized using the Trimmed Mean of M-values (TMM) method^85^, and transformed into log-count-per-million (logCPM) using the voom method^86^. Adjusted RPKM values (Reads Per Kilobase of exon Model per million mapped reads^87^) were computed based on logCPMs and gene model lengths. For two given samples, the RPKM values of the genes was compared using *t*-tests, and a gene was identified as differentially expressed if the corresponding *P* value was less than or equal to 0.05 and the fold-change was greater than 2 at each time point. Only genes with relevant expression levels (RPKM > the square root of the mean expression value of the whole dataset) were considered.

### Alternative splicing analysis

Alternative splicing events were identified as described previously^88^ using the RACKJ software. Three types of alternative splicing were considered, IR, DA, and ES. For detecting IR events, the IR ratio was computed as the average read depth of its intron divided by the average read depth of the neighboring exons, and the IR ratios of three -Fe replicates were compared to those of the controls (+Fe) using *t*-test. Similarly, to detect alternative donor/acceptor or exon skipping events, signals representing AS events (read counts skipping exons, and read counts covering the same splicing junctions) were divided by gene expression levels as background. *T*-tests were performed on the obtained ratios to compare samples from treated plants against control samples. Changes of relative expression levels of AS events were inferred using a *t*-test P value < 0.05 with a fold-change > 2.

### RT-qPCR

Samples were frozen in liquid nitrogen at the end of the experimental period and stored at - 80°C. Total RNA was extracted using the RNeasy Mini Kit (Qiagen) and treated with DNase using the TURBO DNA-free kit (Ambion). Three μg of total RNA per sample was used for obtaining cDNAs. First-strand cDNA was synthesized using oligo(dT) primer and the SupersCript^TM^ III First-Strand Synthesis System (Invitrogen, Cat. No. 18080) for RT-qPCR. The resulting single-stranded complementary cDNAs were then used as a template in real-time RT-PCR assay. RT-qPCRs were carried out with gene-specific primers listed in Table S6, and SYBR^TM^ Green PCR Master Mix (Applied Biosystems, Cat. No. 4367659) according to the manufacturer’s instructions using a QuantStudio 12K Flex Real-Time PCR System.

Three independent replicates were performed for each sample. The ΔΔC_T_ method was used to determine the relative gene expression^89^, with the expression of elongation factor 1 alpha (EF1α; At5g60390) used as an internal control.

### Genomic analyses

Five’ and 3’ splicing site strength scores and event information were downloaded from the PastDB dabase^30^. An inhouse Perl script was developed to associate AS events of PastDB with the accession number of genes from the TAIR10 annotation. The information on transcription factor binding sites and promoter lengths was downloaded from the *Arabidopsis thaliana cis*-regulatory database (AtcisDB) database on the Arabidopsis Gene Regulatory Information Server (AGRIS)^90^.

### UHPLC-MS analysis

Approximately 100 mg plant tissues were harvested, extracted in 1.5 ml of a solution of 375 μl dH_2_O, 750 μl methanol, 375 μl chloroform, and an internal standard (Citrate-2,2,4,4-d_4_, CDN ISOTOPES, Cat. No. 147664-83-3) was added to final concentration of 10 μM for each sample. The supernatant was separated by quick spin-down at 3,000xg, incubated at -20 °C for 30 min, and centrifugated at 3,000xg for 10 min at 4 °C. Obtained supernatant was mixed by vortexing with 375 μl chloroform of chloroform to remove pigments. Colorless supernatant was dried in a SpeedVac, resuspended in 50% methanol, and kept under -80 °C. A Vanquish™ Horizon UHPLC System (Thermo Scientific) coupled to an Orbitrap Fusion Lumos (Thermo Scientific) mass spectrometer was used for the LC-MS analysis. The chromatographic separation for samples was carried out on Atlantis Premier BEH C18 AX VanGuard FIT Column, 1.7 µm, 2.1 x 100 mm column (Waters). The column was maintained at a temperature of 30°C and 1 μL sample were injected per run. The mobile phase A was 2% Acetonitrile 0.1% v/v formic acid in water and mobile phase B was 40% v/v acetonitrile with 20 mM ammonium formate pH 3.0. The gradient elution with a flow rate 0.4 mL/min was performed with a total analysis time of 11 min. The gradient included 0.5% B at 0cmin, a hold at 0.5% B until 2 min, 99.5% B at 6 min, a hold at 99.5% B until 8 min, 0.5% B at 8.5 min, and a hold at 0.5% B until 11 min. General instrumental conditions were RF lens 60%; sheath gas, auxiliary gas, and sweep gas of 50, 10, and 3 arbitrary units, respectively; ion transfer tube temperature of 325 °C; vaporizer temperature of 350 °C; and spray voltage of 3500 V for negative mode. For analysis, a full MS scan mode with a scan range *m*/*z* 50 to 400, resolution 30,000, AGC target 4e5 and a maximum injection time 50 ms was applied. The Xcalibur 4.1 software (Thermo Scientific) was used for the data processing.

## Acknowledgements

We thank Dr. Yuki Nakamura (IPMB, Academia Sinica) for kindly providing chemicals for UHPLC standard and Shou-Jen Chou in the Genomic Technology Core Facility at IPMB for preparing libraries for RNA-seq analysis. We also thank Mei-Jane Fang at the Genomic Technology Core Facility at the Institute of Plant and Microbial Biology for using of the QuantStudio 12K Flex Real-Time PCR System, and Yu-Ching Wu for the UHPLC-MS analysis, which was supported by the Academia Sinica Metabolomics Core Facility at the Agricultural Biotechnology Research Center of Academia Sinica, supported by Academia Sinica Core Facility and Innovative Instrument Project (AS-CFII-108-108).

## Author contributions

E.J.H. and W.S. designed the research; E.J.H. performed the experiments; E.J.H., W.D.L., and W.S. analysed the data; W.S. and E.J.H., wrote the manuscript.

## Conflict of interest

The authors declare no conflict of interest.

## Data availability

The RNA-seq data have been deposited at NCBI under the accession number PRJNA759647 Reviewer link: https://dataview.ncbi.nlm.nih.gov/object/PRJNA759647?reviewer=itt5n0pfr6mmqho1akj54r5g2k

## Notes

### Competing Interest Statement

The authors have declared no competing interest.

