## Supplementary Figures S1-S6 for "Genomically hardwired regulation of gene activity orchestrates cellular iron homeostasis in Arabidopsis"

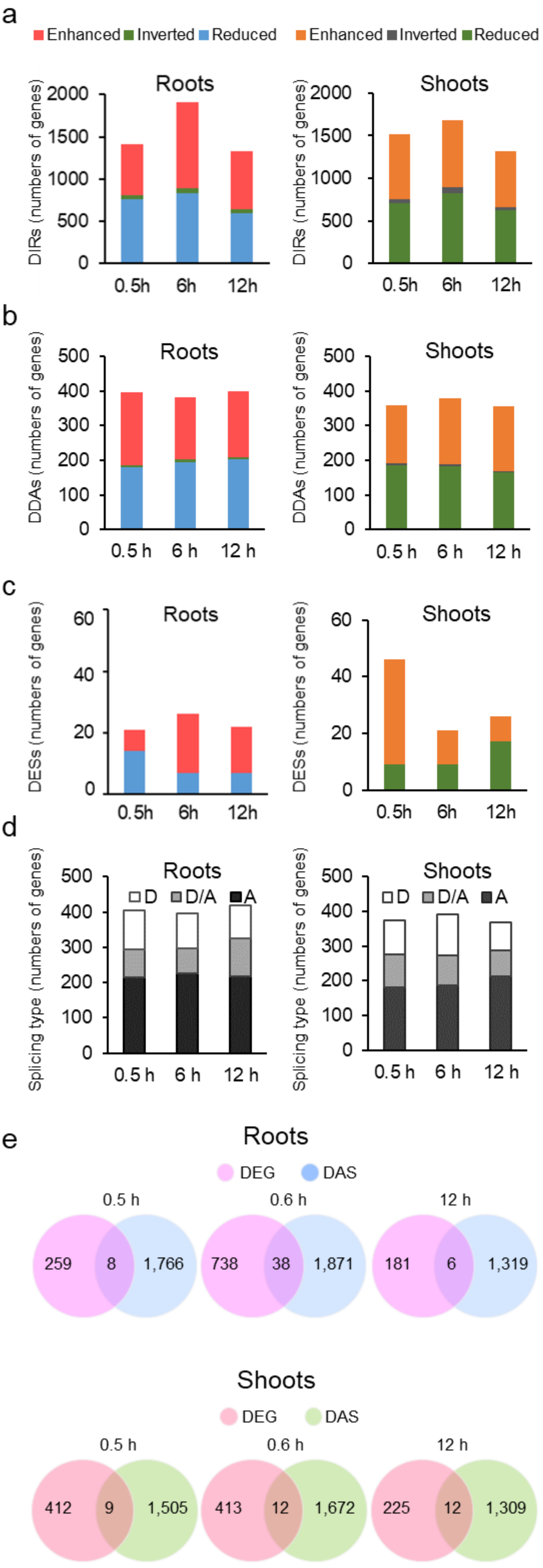


**Supplementary Figure S1.** Regulation of differential alternative splicing events (DAS) and overlap with differentially expressed genes (DEGs). a) Regulation of differential intron retention (DIR) events. b) Regulation of differential alternative donor/acceptor (DDA) events. c) Regulation of differential exon skipping (DES) events. d) Separation of alternative donor (D), alternative donor/acceptor (D/A), and alternative acceptor (A) events within the DDA genes. e) Venn diagrams indicating the overlap of DEG and DAS at the various timepoints in roots (upper panel) and shoots (lower panel). 'Enhanced' and 'reduced' refers to increased and decreased fold changes of alternative splicing features, respectively. 'Inverted' denotes genes that harbour both enhanced and reduced features.

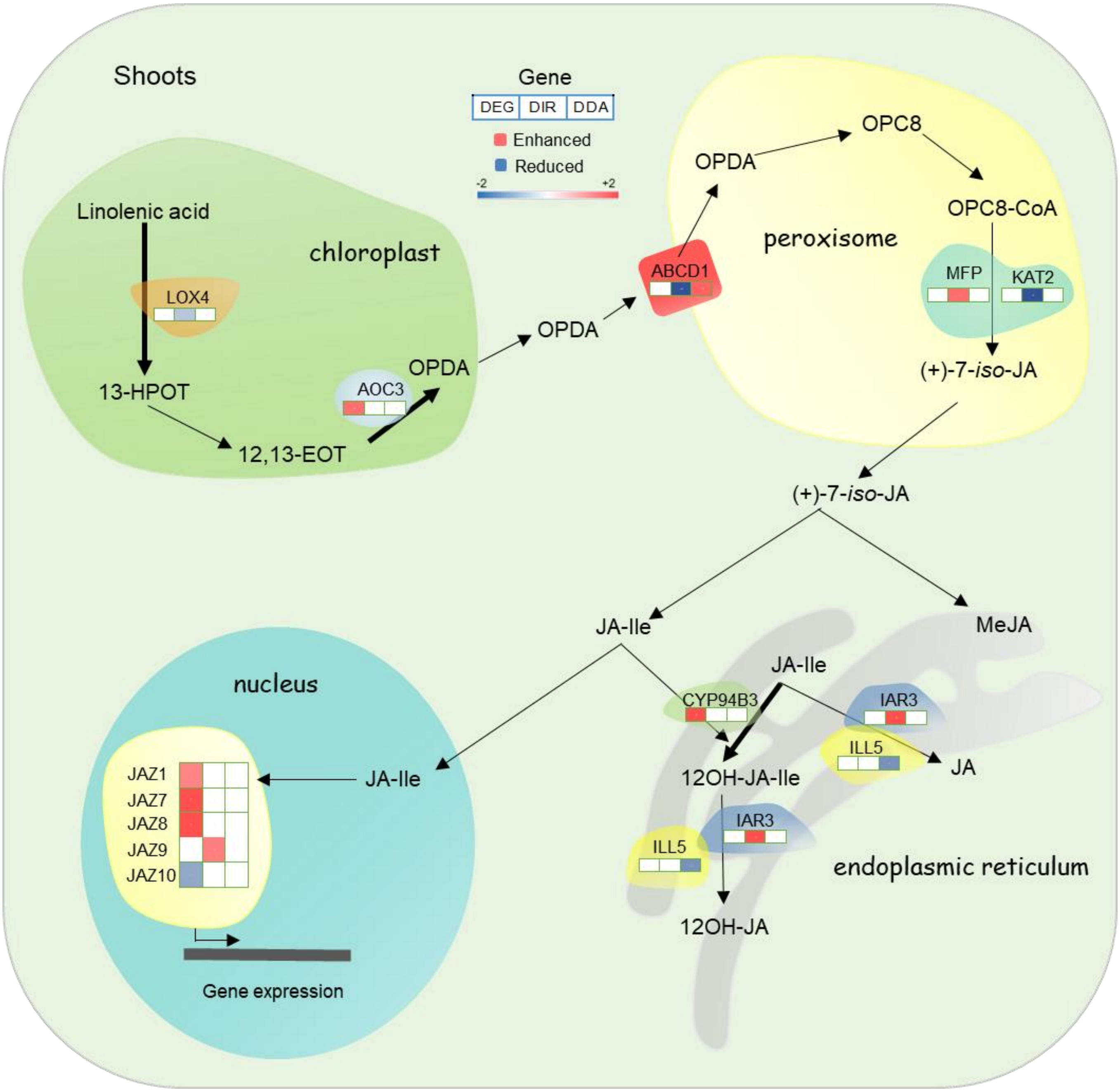


**Supplementary Figure S2.** Induction of genes involved in jasmonate biosynthesis, catabolism, and signalling in shoots of plants exposed to short-term Fe deficiency treatment. In plastids, lipid-derived α-linolenic acid is converted by LOX and AOC into oxophytodienoic acid (OPDA). In the cytoplasm, JA is conjugated with amino acids to JA-Ile by JAR1 or to MeJA by JMT. In the endoplasmic reticulum, JA-Ile is degraded to 12OH-JA-Ile and converted to 12-OH-JA via IAR3, ILL5. The latter enzymes can also convert JA-Ile to JA. In the absence of nuclear JA-Ile, expression of JA responsive genes via MYC2 is repressed by JAZ. Upon the entry of JA-Ile into the nucleus, JAZ is degraded via the 26S proteasome and ultimately triggers transcriptional activation of the target genes. Only genes that were differentially expressed or harbored DAS features in response to Fe starvation are shown.

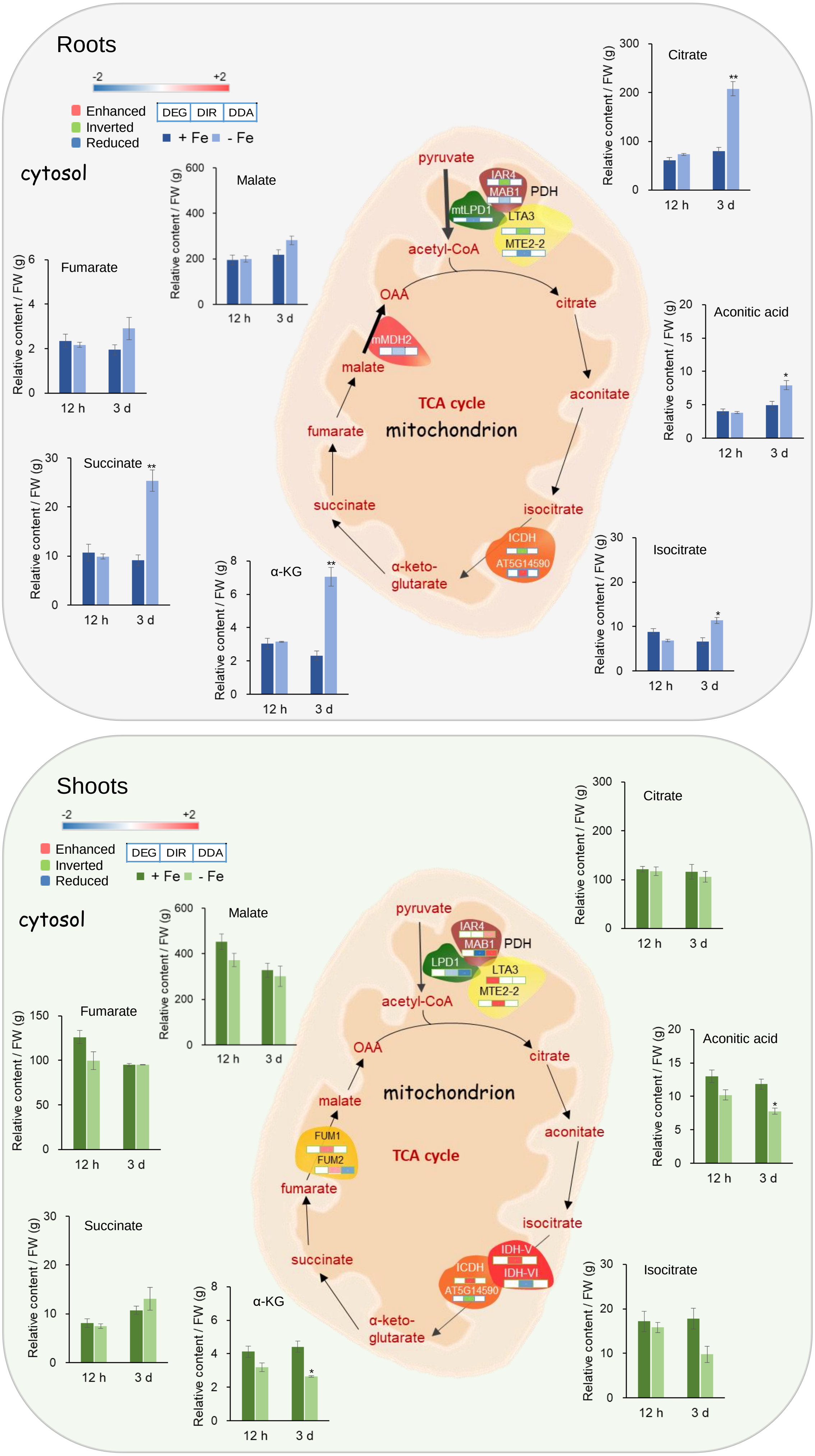

Roots
-2
+2
Enhanced
Inverted
Reduced
DEG
DIR
DDA
+ Fe
- Fe
cytosol
Fumarate
Malate
Succinate
α-KG
Citrate
Aconitic acid
Isocitrate
Relative content / FW (g)
12 h
3 d
pyruvate
IAR4
MAB1
PDH
mtLPD1
LTA3
MTE2-2
acetyl-CoA
OAA
citrate
mMDH2
malate
aconitate
TCA cycle
fumarate
mitochondrion
succinate
isocitrate
ICDH
AT5G14590
α-keto-
glutarate
Shoots
-2
+2
Enhanced
Inverted
Reduced
DEG
DIR
DDA
+ Fe
- Fe
cytosol
Malate
Fumarate
Succinate
α-KG
Citrate
Aconitic acid
Isocitrate
pyruvate
IAR4
MAB1
PDH
LPD1
LTA3
MTE2-2
acetyl-CoA
OAA
citrate
malate
mitochondrion
aconitate
FUM1
FUM2
TCA cycle
fumarate
isocitrate
IDH-V
IDH-VI
ICDH
AT5G14590
succinate
α-keto-
glutarate

**Supplementary Figure S3.** Regulation of enzymes of the TCA cycle and their products by Fe-deficiency in roots (upper panel) and shoots (lower panel). 'Enhanced' and 'reduced' refers to increased and decreased fold changes of alternative splicing features, respectively. 'Inverted' denotes genes that harbour both enhanced and reduced features. TCA intermediates were determined 12 hours and 3 days after transfer to Fe-deplete media.

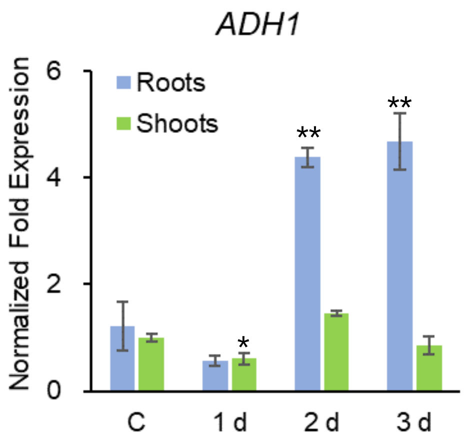


**Supplementary Figure S4.** RT-qPCR analysis of *ADH1* in response to Fe deficiency. Gene expression was determined by RT-qPCR using $\Delta\Delta C_T$ method and expression of elongation factor 1 alpha as an internal control. Each bar represents the mean ± SE of three independent experiments. Asterisks indicate significant differences from the wild type in each treatment: *, $P \leq 0.05$; **, $P \leq 0.01$; ***, $P \leq 0.001$. C, control.

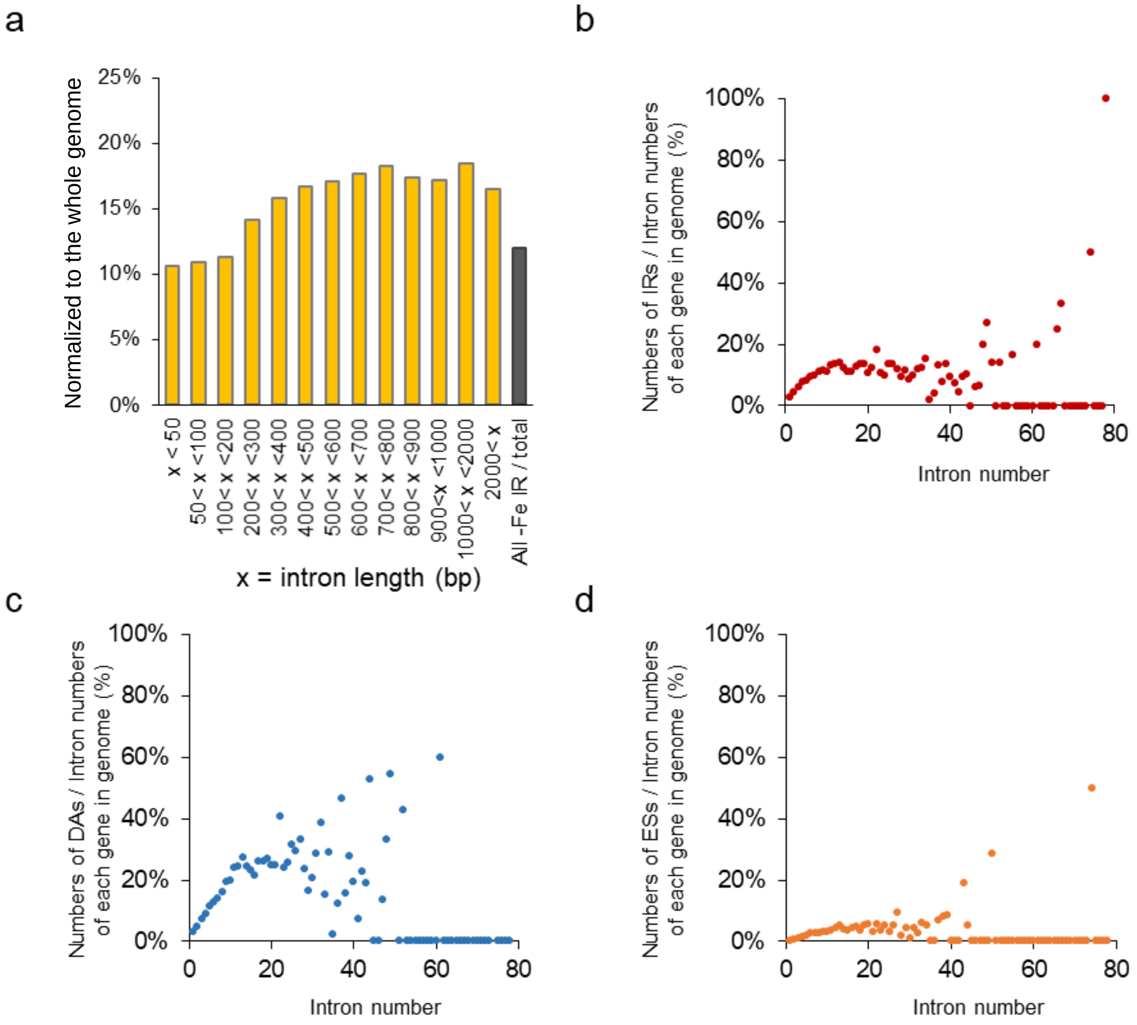


**Supplementary Figure S5.** Effects of intron length and number on alternative splicing. a) Effect of intron length on IR. b) Effect of intron number on IR. c) Effect of intron number on DA. d) Effect of intron number on ES.

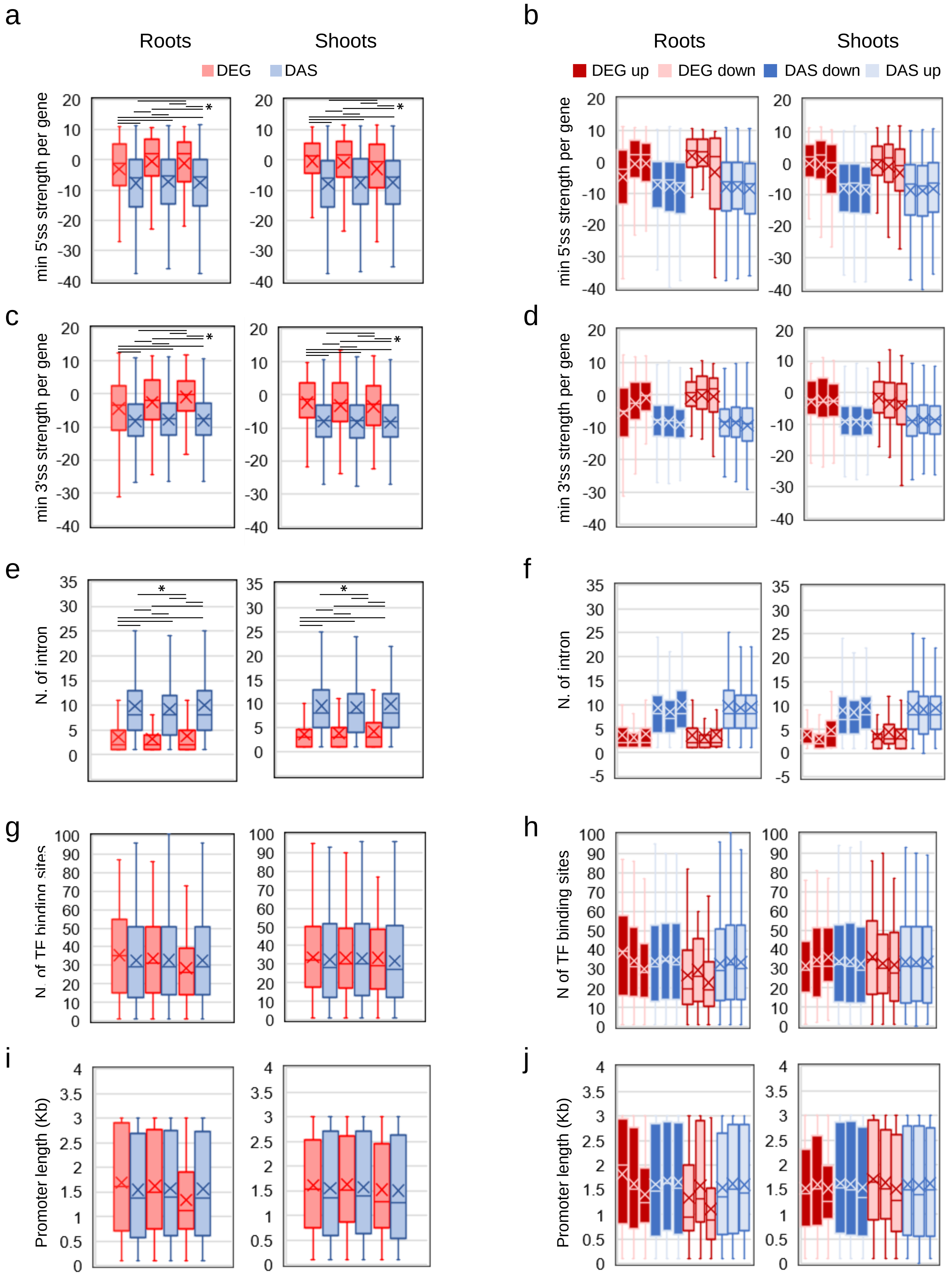


**Supplementary Figure S6.** Effects of genomic features on the mode of gene regulation after 0.5, 6, and 12 hours exposure to Fe deficiency (left to right). a) Minimum 5' splice site strength. b) Effect of minimum 5' splice site strength on DEG and DAS in up- and downregulated genes. c) Minimum 3' splice site strength. d) Effect of minimum 3' splice site strength on DEG and DAS in up- and downregulated genes. e) Number of transcription factor binding sites (TFBS). f) Effect of TFBS on DEG and DAS in up- and downregulated genes. g) Promoter length. h) Effect of promoter length on DEG and DAS in up- and downregulated genes.
